# Multi-immersion open-top light-sheet microscope for high-throughput imaging of cleared tissues

**DOI:** 10.1101/548107

**Authors:** Adam K. Glaser, Nicholas P. Reder, Ye Chen, Chengbo Yin, Linpeng Wei, Soyoung Kang, Lindsey A. Barner, Weisi Xie, Erin F. McCarty, Chenyi Mao, Aaron R. Halpern, Caleb R. Stoltzfus, Jonathan S. Daniels, Michael Y. Gerner, Philip R. Nicovich, Joshua C. Vaughan, Lawrence D. True, Jonathan T.C. Liu

**Affiliations:** Department of Mechanical Engineering, University of Washington, Seattle, WA USA; Department of Pathology, University of Washington, Seattle, WA USA; Department of Chemistry, University of Washington, Seattle, WA USA; Department of Immunology, University of Washington, Seattle, WA USA; Applied Scientific Instrumentation, Eugene, OR USA; Allen Institute for Brain Science, Seattle, WA USA; Department of Physiology and Biophysics, University of Washington, Seattle, WA USA

## Abstract

Recent advances in optical clearing and light-sheet microscopy have provided unprecedented access to structural and molecular information from intact tissues. However, current light-sheet microscopes have imposed constraints on the size, shape, number of specimens, and compatibility with various clearing protocols. Here we present a multi-immersion open-top light-sheet microscope that enables simple mounting of multiple specimens processed with a variety of protocols, which will facilitate wider adoption by preclinical researchers and clinical laboratories.

**One Sentence Summary:** Glaser *et al.* describe a multi-immersion open-top light-sheet microscope that enables simple and high-throughput imaging of large numbers of preclinical and clinical specimens prepared with a variety of clearing protocols.

## Main Text

Recent advances in tissue clearing have provided unprecedented visual access to structural features and molecular targets within large intact specimens [1]. These clearing approaches have the potential to accelerate new discoveries across multiple fields of research, including neuroscience, developmental biology, and anatomic pathology [2–5] (**Supplementary Figure 1**). However, fully harnessing the benefits of tissue clearing requires user-friendly and versatile microscopes for three-dimensional (3D) imaging of a wide variety of content-rich specimens.

Over the past decade, light-sheet fluorescence microscopy (LSFM) has emerged as the technique of choice for fast and gentle 3D microscopy of relatively transparent specimens [6, 7]. In LSFM, a thin sheet of light illuminates a specimen such that fluorescence is selectively excited within a single “optical section”. The fluorescence is imaged onto a high-speed camera in the direction perpendicular to the light sheet. Initial LSFM systems were purposefully designed for imaging small model organisms, often living, over unprecedented long timescales [8]. However, in recent years, the advantages of LSFM have also been harnessed for 3D imaging of large cleared tissues, where speed is the main advantage over alternative microscopy methods [9–11]. Unfortunately, most LSFM architectures impose constraints on the size, shape, number of specimens, and/or compatibility with various tissue clearing protocols (**Supplementary Discussion**). In addition, specimen mounting has typically been complex and slow, which has hampered the application of LSFM for high-throughput imaging (**Supplementary Figure 2**).

Here, we have developed an easy-to-use multi-immersion open-top light-sheet (OTLS) microscope in which all optical components are positioned below a transparent holder so that specimens may be conveniently placed on top in a manner similar to a “flatbed scanner” (**Fig. 1a** and **Supplementary Video 1**) [5, 12, 13]. An air objective delivers an illumination light sheet through an immersion chamber, holder, and into a specimen, where fluorescence is generated and then collected by a multi-immersion objective in the direction orthogonal to the light sheet. The multi-immersion capability of our OTLS system spans the refractive index range of all published tissue-clearing methods (including expansion, aqueous, and solvent-based protocols). A key to achieving this is by precisely matching the refractive index of the immersion medium, holder material, and specimen (**Fig. 1b**). A plano-convex lens with a radius of curvature matched to the focus of the illumination beam acts as a solid immersion lens (SIL) to prevent spherical and off-axis aberrations from occurring as the light sheet enters the system. The point spread function (PSF) of the system is shown in **Fig. 1c**, in which sub-micron resolution is achieved lateral to the collection axis. Resolution along the collection axis (~3.5 μm) is comparable to the thickness of a typical slide-mounted histology section. Note that the numerical apertures (NA) of both the illumination and collection beams are proportional to the matched refractive index of the immersion medium, holder material, and specimen (**Fig. 1d**). The open-top architecture (**Fig. 1e**) enables fast and simple mounting of multiple specimens with diverse shapes such as human biopsies, thin tissue slices, and whole mouse organs in modular holders (**Fig. 1f**). Large-volume imaging at a speed of ~1 min/mm^3^ is achieved entirely through stage scanning, in which a series of adjacent volumetric image strips (scanned in the *x* dimension) are tiled in the lateral (*y*) and vertical (*z*) directions (**Fig. 1g** and **Supplementary Video 2**). The working distance of the collection objective allows for an imaging depth of up to 0.5 cm. In comparison to a previously published open-top prototype [5], our new OTLS system exhibits (1) improved axial and lateral resolution (an order-of-magnitude reduction in the focal volume), (2) ~20× greater imaging depth, (3) mitigation of shadowing artifacts, and (4) multi-immersion capabilities (**Supplementary Figure 3**).

**Figure 1.**
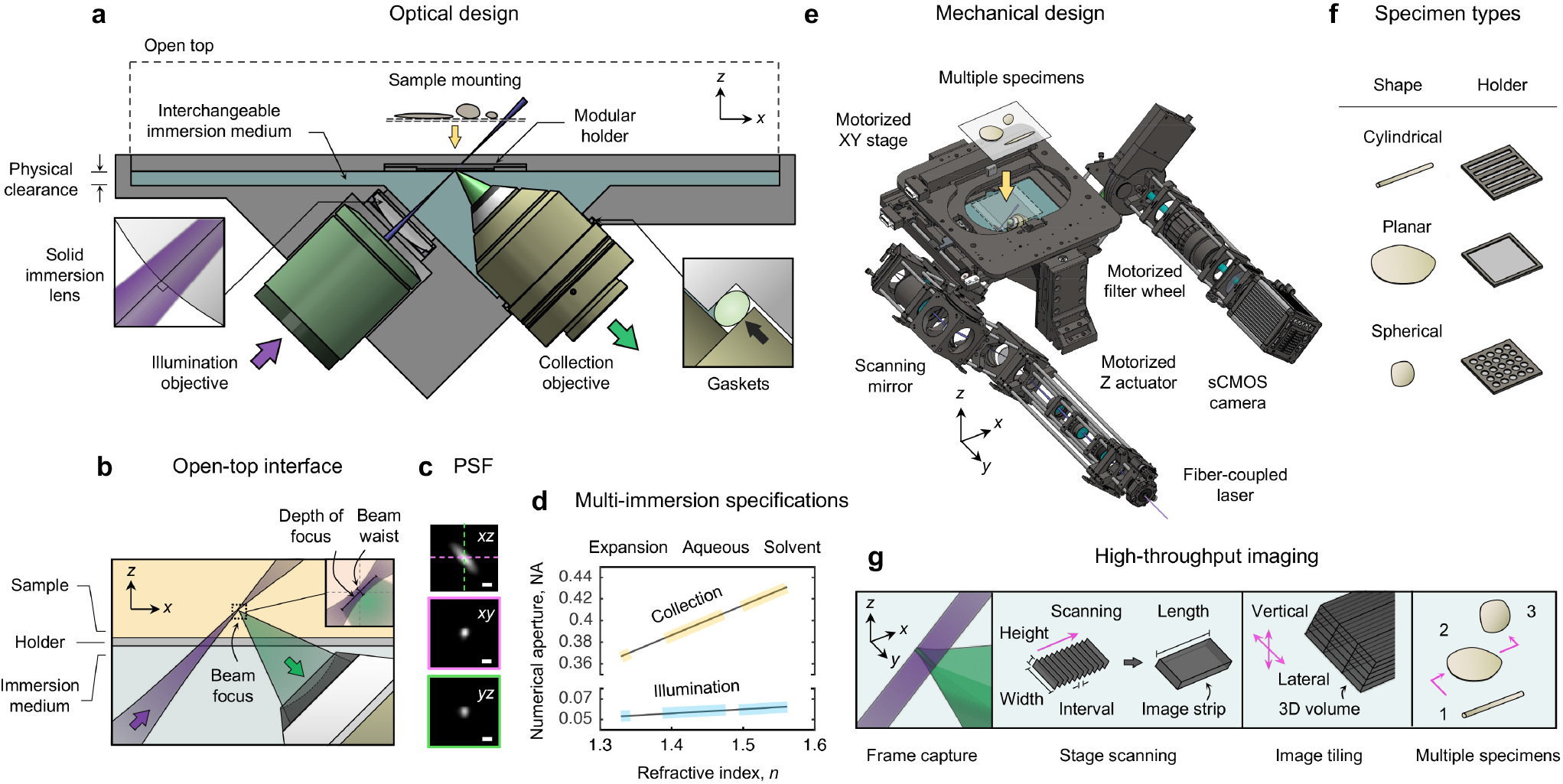
Multi-immersion open-top light-sheet (OTLS) microscope. **(a)** The system enables simple mounting of multiple specimens with modular transparent holders. Illumination and collection objectives are located underneath the specimen holders and are separated by a liquid reservoir filled with an interchangeable immersion medium. **(b)** The off-axis illumination light sheet and collected fluorescence travel obliquely through the immersion media, holder, and specimen. Aberrations are minimized by precisely matching the refractive index of all three materials, and by utilizing the wavefont-matching properties of a solid immersion lens (SIL) along the illumination path. The depth-of-focus and beam waist of the light sheet are depicted in the inset (upper right). The point spread function (PSF) of the system and refractive-index-dependent numerical aperture (NA) of the illumination and collection beams are shown in **(c)** and **(d)**. **(e)** The mechanical design of the system includes a motorized XY stage, motorized Z actuators, motorized filter wheel, scanning mirror, computer-controlled multi-wavelength fibre-coupled laser package, and sCMOS camera, all of which enable high-throughput automated imaging of multiple specimens simply placed on a flat plate, or placed within a diverse assortment of transparent holder designs **(f)**. **(g)** Volumetric imaging is achieved by using a combination of stage-scanning and lateral/vertical tiling. The scale bars in **(c)** denote 1 μm.

Since precise index matching of the immersion media, holder, and specimen are necessary for aberration-free imaging, we performed optical simulations (**Fig. 2a**) to explore the tolerance of the system to the optical path difference (Δ*n*×*t*) introduced by a holder with a refractive index mismatch, Δ*n*, and thickness, *t*. We quantified the Strehl Ratio, *S*, of the system as a function of Δ*n*×*t*, and determined that Δ*n*×*t* < 0.002 is necessary for near-diffraction-limited imaging (*S* > 0.8). Based on these findings, we surveyed glasses and monomers/polymers as potential holder materials, and determined the maximum-allowed thickness, *t*_*max*_, based upon the intrinsic mismatch, Δ*n*, between these materials and published clearing protocols (**Fig. 2b**). These chemical-reagent and material combinations are enabling for our OTLS system, and are beneficial for other cleared tissue imaging systems. To validate the multi-immersion capabilities of our system, we fabricated customized holders for a variety of clearing protocols spanning solvent, aqueous, and expansion-based protocols, and performed high-throughput imaging of diverse tissue types and shapes (a summary of the imaging parameters for all specimens are shown in **Supplementary Table 3**).

**Figure 2.**
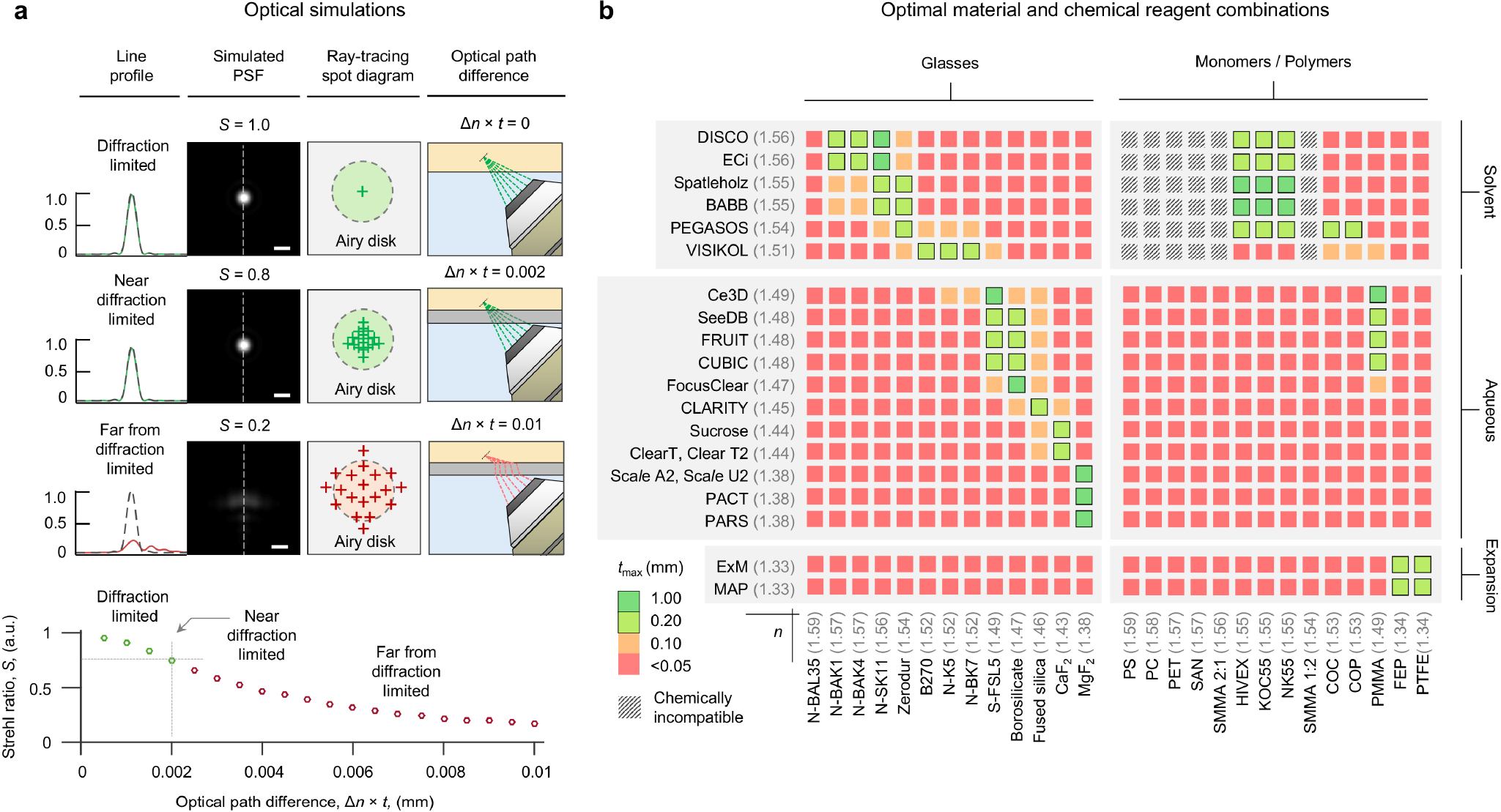
Holder design for OTLS imaging. **(a)** Optical simulations of the system’s PSF, and ray-tracing spot diagrams, are shown for scenarios in which the optical path difference (Δ*n*×*t*) is negligible, small, or large, which results in diffraction-limited (*S* ~ 1.0), near-diffraction-limited (*S* > 0.8), or aberrated (*S* < 0.8) imaging performance, respectively. The dependence of the Strehl Ratio, *S*, as a function of Δ*n*×*t* is plotted, indicating that for diffraction-limited imaging, the condition that Δ*n*×*t* < 0.002 should be maintained. Based on this condition, potential glass and monomer/polymer holder materials are shown in **(b)**. The color scale indicates the maximum material thickness, *t*_*max*_, that is allowed based upon the intrinsic mismatch, Δ*n*, of those materials with published clearing protocols. Chemically incompatible combinations of materials and chemical reagents are also indicated.

Solvent-based protocols involve dehydration of tissue specimens and replace the water with organic reagents of a higher refractive index (*n* = 1.51 − 1.56). These solvents are optically compatible with several high-index glasses. Unfortunately, these higher-index glasses also have high Abbe numbers (i.e. a low variation in refractive index versus wavelength) compared to organic solvents, which typically have low Abbe numbers. In addition, glasses, which are brittle, must be relatively thick and are not easily machined. Therefore, we explored the use of several monomers/polymers. Despite being optically compatible, we observed that styrene-based polymers (e.g. PS, SMMA, and SAN) were all destroyed after exposure to solvent-based clearing reagents. However, we identified three optically and chemically compatible resin-based monomers (HIVEX, NK55, and KOC55, used for manufacturing eyeglass lenses) that are ideal for many solvent-based clearing protocols, including DISCO, BABB, and ECi [14–18]. We demonstrated the potential clinical utility of our system by imaging multiple human prostate biopsies *in toto* (**Fig. 3a** and **Supplementary Videos 3 and 4**). We found that ECi-clearing (*n* = 1.56) is well-suited for this application due to its clearing efficacy, as well as low toxicity [19]. A custom HIVEX holder (*n* = 1.55) was machined with channels to accommodate multiple biopsies (13 in this case) and to keep them aligned and parallel. Due to the effective tissue clearing of ECi and precise refractive index matching with the HIVEX holder, we were able to resolve sub-nuclear features in benign and malignant prostate glands throughout the entire 1-mm diameter of all biopsies. As noted by us and others, this ability to visualize the 3D structure of human cancers should improve prognostication and treatment decisions [3, 5, 20].

**Figure 3.**
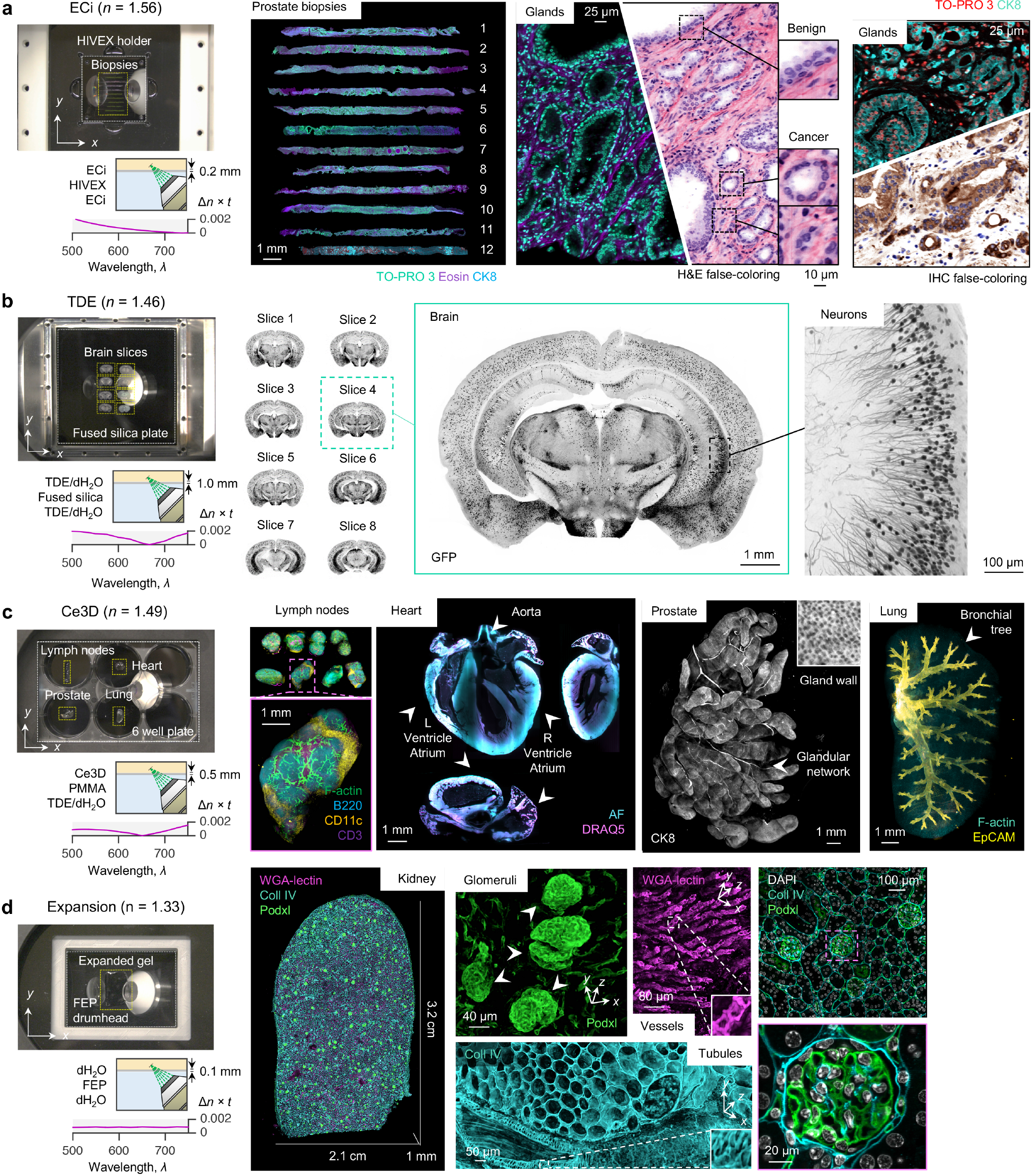
Multi-immersion and multi-sample imaging. **(a)** OTLS imaging of 12 ECi-cleared human prostate biopsies placed within a multi-biopsy holder. Zoomed-in views illustrate the complex 3D structure of benign and malignant glands. Images can be false-colored to mimic the appearance of conventional chromogen-based (absorption-based) H&E and IHC histopathology. **(b)** High-throughput imaging of 8 TDE-cleared mouse brain slices placed on a 10- by 10-cm glass plate. A higher-magnification region of interest demonstrates the ability to visualize individual neurons. **(c)** Whole-organ imaging of Ce3D-cleared mouse lymph nodes, heart, prostate, and lung placed within an multi-well plate. **(d)** Large-scale imaging of an expanded thick-kidney slice placed on a teflon drumhead. High-resolution regions of interest show individual glomeruli, vessels, and tubules.

Aqueous-based protocols typically involve removing lipids using detergents, followed by immersion in diluted water-soluble reagents, resulting in a refractive index range of *n* = 1.38 to 1.49 [21]. For most aqueous-based protocols, there is an optically compatible glass such that Δ*n* ≤ 0.01. For example, fused silica (*n* = 1.46) is well-suited for several protocols, including CLARITY [2, 22]. However, most transparent monomers/polymers (with the exception of PMMA) have either a high (*n* > 1.49) or low (*n* < 1.38) refractive index and are therefore not optically compatible with aqueous-based protocols. To demonstrate compatibility of our system with aqueous protocols, we cleared adjacent slices of a mouse brain (200 μm thick) using simple immersion in diluted TDE (*n* = 1.46) and imaged them using a 1 mm thick fused-silica plate (*n* = 1.46) with a large 10×10 cm viewing area (**Fig. 3b** and **Supplementary Video 5**). High-magnification views of our datasets enabled unambiguous visualization of neuronal structures. We also cleared and labeled multiple mouse organs using the Ce3D protocol (*n* = 1.49) and imaged them in a single automated session using a customized 6-well plate with a 0.5 mm thick PMMA bottom substrate (*n* = 1.49) (**Fig. 3c** and **Supplementary Video 6**). Lymph nodes, heart, prostate, and lung tissues (1 to 4 mm thick) were mounted in separate wells and imaged *in toto* for 3D visualization of immune cells in lymph nodes, the ventricles and valves within a heart, the glandular network within the prostate, and the bronchial tree within the lung [23]. Note that the use of multi-well plates (e.g. 6-well or 96-well plates) prevents contamination between samples, which is particularly important for clinical assays and cell culture (a feature not possible with most other LSFM systems, see **Supplementary Discussion**).

Expansion-based protocols provide a magnified view of structures that are otherwise too small to resolve with a given microscope [24–26]. To date, these expanded hydrogel specimens have consisted mostly of water and therefore have a refractive index close to that of pure water (*n* = 1.33). While there are currently no glasses at this refractive index, fluoro-polymers (i.e., Teflons), including fluorinated ethylene-propylene (FEP) and poly-tetra-fluoro-ethylene (PTFE) possess a compatible refractive index (*n* = 1.34). These materials can be manufactured as thin sheets and stretched tight as “drumhead” surfaces that are ideal holders for expanded specimens. Using a customized drumhead, we imaged a 4× expanded 200 μm thick kidney section (**Fig. 3d** and **Supplementary Videos 7-9**). After expansion, the physical size of the tissue was 2.1×3.2×0.1 cm. Representative zoomed-in views provide a highly detailed view of 3D structures such as glomeruli, renal tubules, and blood vessels.

We have developed and characterized a multi-immersion OTLS microscope that enables high-throughput automated imaging of optically cleared specimens with an ease of use that should facilitate broader adoption of light-sheet-based 3D microscopy by both researchers and clinicians. Our system imposes minimal constraints on specimen shape/size and allows for fast and convenient mounting of multiple tissue specimens for automated imaging. The system provides an imaging depth of 0.5 cm over a maximum lateral area of 10×10 cm at a speed of ~ 1 min/mm^3^, which can be tailored for specific research applications in future designs (see **Supplementary Figure 22**). We have shown that our system can interface with a variety of modular specimen holders tailored for specific tissue types and clearing protocols, with the ability to isolate different specimens in individual wells. Due to its open-top geometry, our system also provides unsurpassed versatility to interface with future tissue-based protocols and a wide range of potential accessory technologies such as microfluidic devices, single-cell electrophysiology, and micro-aspiration [27–30].

## Online Methods

### Multi-immersion open-top light-sheet microscope

An optical schematic of the system is shown in **Supplementary Figure 4** and was modeled using commercially available ray-tracing software (ZEMAX LLC) (**Supplementary Figs. 5** and **6**, available as **Supplementary ZEMAX Files**). Illumination light is coupled into the system by a single-mode fiber with a numerical aperture of 0.12 from a four-channel digitally controlled laser package (Skyra, Cobolt Lasers). Light emanating from the fiber is collimated with a lens, L1 (AC1-128-019-A, *f* = 19 mm), and then expanded along one axis using a 3× cylindrical telescope consisting of lenses, C1(ACY-254-50-A, Thorlabs, *f* = 50 mm) and C2 (ACY-254-150-A, Thorlabs, *f* = 150 mm) to provide multi-directional illumination (**Supplementary Figure 7**) [1]. The resulting elliptical Gaussian beam is then relayed to the scanning galvanometer, GM (6210H, Cambridge Technology) using lenses R1 (AC-254-100-A, Thorlabs, *f* = 100 mm) and R2 (AC-254-050-A, Thorlabs, *f* = 50 mm). The scanning mirror is driven by a sinusoidal voltage from a waveform generator (PCI-6115, National Instruments) at a frequency of 800 Hz. The scanning beam is relayed to the back focal plan of the illumination objective (XLFLUOR340/4× 0.28 NA, Olympus) using a scan lens, SL (CLS-SL, Thorlabs, *f* = 70 mm) and tube lens, TL1 (TTL200, Thorlabs, *f* = 200 mm). Finally, the elliptical beam travels through a plano-convex lens (LA4725, Thorlabs, *R* = 34.5 mm), immersion medium, holder, and finally specimen.

Fluorescence is collected by a multi-immersion objective (#54-10-12, Special Optics, distributed by Applied Scientific Instrumentation). This provides <1 μm in-plane resolution for all immersion media (**Supplementary Figure 8**). The fluorescence is filtered with a motorized filter wheel (FW102C, Thorlabs) with band-pass filters for the 405 nm (FF02-447/60-25, Semrock), 488 nm (FF03-525/50-25, Semrock), 561 nm (FF01-618/50-25, Semrock), and 638 nm (FF01-721/65-25, Semrock) excitation wavelengths. The filtered fluorescence is focused onto a 2048×2048 pixel sCMOS camera (ORCA-Flash4.0 V2, Hamamatsu) by a tube lens, TL2 (TTL165, Thorlabs, *f* = 165 mm). The tube lens provides a Nyquist sampling of ~0.45 μm/pixel, which provides a horizontal field of view of ~0.9 mm over the 2048 pixels of the camera. The vertical field of view is reduced to 256 pixels to match the depth of focus of the illumination light sheet (~110 μm). The 256 pixels are oriented parallel to the rolling shutter readout direction of the camera, which provides an exposure time of 1.25 ms and a framerate of 800 Hz. The maximum imaging depth is limited by the physical clearance of the holder and collection objective (0.5 cm). The illumination objective, solid immersion lens, and collection objective interface with the immersion chamber through customized aluminum mounts (**Supplementary Figure 9**) which are available as **Supplementary CAD Files**.

Image strips are collected with a combination of stage-scanning and lateral/vertical tiling using a motorized XY stage and Z actuators (FTP-2050-XYZ, Applied Scientific Instrumentation - ASI). The stage-scanning firmware is used to send a TTL trigger signal from the XY stage to the sCMOS camera for reproducible start positioning (<1 μm) of each image strip (**Supplementary Figure 10**). The spatial interval between successive frames is set to ~0.32 μm, which, given the 800 Hz camera framerate, corresponds to a constant stage velocity of ~0.25 mm/sec. For lateral tiling, an offset of 0.8 mm between adjacent image strips is used (~11% overlap). For vertical tiling, the 110 μm depth of focus is oriented at 45 deg., which corresponds to an image strip height of ~80 μm. Therefore, a vertical tiling offset of 70 μm is used (~12% overlap). The laser power is increased with depth per a user defined attenuation coefficient, *P* = *P*_*0*_ ×*exp*(*z*/*μ*), to account for the attenuation of the illumination light sheet as it penetrates deeper into the specimen. The entire image acquisition is controlled by a custom LabVIEW (National Instruments) program. As shown in **Supplementary Figure 11**, the program consists of a series of nested loops for imaging multiple specimens, collecting multiple color channels, and lateral/vertical tiling. A complete list of components is available in **Supplementary Table 1**.

### Computer hardware

During acquisition, the images are collected at the maximum data-transfer rate (~800 MB/sec) by a dedicated workstation (Precision Tower 5810, Dell) equipped with a CameraLink interface (Firebird PCI Express, Active Silicon). The data is streamed in real-time using the proprietary DCIMG Hamamatsu format to a mapped network drive located on an in-lab server (X11-DPG-QT, SuperMicro) running 64-bit Windows Server, equipped with 384 GB RAM and TitanXP (NVIDIA) and Quadro P6000 (NVIDIA) GPUs. The server contains two high-speed RAID0 storage arrays of 4×2.0 TB SSDs, as well as a larger direct-attached RAID6 storage array with 15×8.0 TB HDDs. All RAID arrays are hardware-based, the RAID0 arrays are controlled by an internal 8-port controller (LSI MegaRaid 9361-8i 1 GB cache) and the RAID6 array is controlled by an external 8-port controller (LSI MegaRaid 9380-8e 1 GB cache). Both the server and acquisition workstation are equipped with 10G SFP+ network cards, jumbo frames, and parallel send/receive processes matched to the number of computing cores on the workstation (8 physical cores) and server (16 physical cores), which reliably enables >1.0 GB/sec network transfer speeds (to accommodate the data-transfer rate of the sCMOS camera and enable simultaneous data-processing routines). The hardware setup is shown in **Supplementary Figure 12**. The complete hardware configuration is listed in **Supplementary Table 2**.

### Data processing and visualization

Collected datasets undergo a Python pre-processing routine before being visualized in 2D and 3D by several open-source and commercial packages. Each image strip is stored in a single DCIMG file. These DCIMG files are read into RAM by a DLL compiled using the Hamamatsu DCIMG software development kit (SDK) and first de-skewed at 45 deg. By precisely setting the interval between successive frames, the de-skewing is quickly performed by simply shifting each plane of pixels in the image strip by an integer pixel offset (**Supplementary Figure 13**). This operation is extremely fast compared to alternative de-skewing approaches using computationally expensive affine transformations. The data is then written from RAM to disk using the Hierarchical Data Format (HDF5) with the metadata and XML file structured for subsequent analysis using BigStitcher [2]. A custom HDF5 compression filter (B3D) is used with default parameters to provide ~10× compression which is within the noise limit of the sCMOS camera [3]. This pre-processing routine is applied to all DCIMG files, ultimately resulting in a single HDF5/XML file for BigStitcher. The alignment of all image strips is performed in BigStitcher, and finally fused to disk in either TIFF or HDF5 file formats. The resulting TIFF and HDF5 files are then visualized using open-source and commercial packages, including ImageJ, BigDataViewer, Aivia (DRVision), and Imaris (Bitplane) [4, 5]. To optionally provide false-colored pseudo-H&E histology images, a Beer-Lambert coloring algorithm is applied using a Python script [6]. The entire processing pipeline is shown in **Supplementary Figure 14** and available as **Supplementary Code**.

### Specimen holders

All holders were attached to the motorized XY stage using custom machined aluminum adapters plates (HILLTOP21). For the mouse brain slices, a 1-mm thick fused silica window (Esco Optics) with a 10×10-cm cross-section was attached to a custom adapter plate using UV-curing glue (**Supplementary Figure 15**). Mouse organs cleared using Ce3D were imaged on a customized 6-well plate. The bottom of a conventional polystyrene 6-well plate (Cat:CLS3506, Sigma-Aldrich) was removed and replaced with a 0.5 mm thick PMMA plate (Goodfellow USA) (**Supplementary Figure 16**). For the expanded kidney specimen, a custom “drumhead” was fabricated and adapted for mounting to the microscope. The drumhead tightens a 0.1 mm thick FEP film over an extruded opening, which is ideal for OTLS imaging of expanded specimens (**Supplementary Figure 17**). To overcome the hydrophobic nature of the FEP films (which cause drifting of expanded specimens), the upper surface of the FEP films were treated with 0.1% (w/v) poly-lysine (Cat:P8920, Sigma-Aldrich) for charged-based adhesion of specimens to the FEP surface. For the human prostate biopsies, HIVEX lens blanks (Conant Optical) were purchased and custom machined using an in-house desktop mill (OtherMill, Bantam Tools) (**Supplementary Figure 18**). The 1/8 inch, 1/16 inch, and 1/32 inch drill bits were used, and the feed rates and drill speeds were optimized for the HIVEX material. CAD files for all sample holders are available as **Supplementary CAD Files**. The system can also be used as a whole-slide scanner for conventional fluorescently labeled histology slides using a commercially available slide holder (MLS203-P2, Thorlabs) (**Supplementary Figure 19**). Dispersion curves for the various holder materials and clearing reagent combinations are shown in **Supplementary Figure 20**.

### Optical simulations

Optical simulations were performed using commercially available ray-tracing software (ZEMAX, LLC) with a “blackbox” model of the multi-immersion objective (provided by the manufacturer, Special Optics). For the simulations shown in **Fig. 2**, the base refractive index of the immersion medium and specimen was assumed to be *n* = 1.45, and the optical path difference was varied. For all scenarios, the imaging depth was set to 1 mm, and the PSF was measured at the center of the imaging field of view. The same relationship between Strehl Ratio and optical path difference was observed for other base refractive-indices and imaging depths, under the assumption that the optical properties of the immersion medium and specimen were the same. The ZEMAX files for the OTLS system are available as **Supplementary ZEMAX Files**.

### Collection and processing of mouse brain slices

A mouse of line Sst-IRES-Cre;Ai139(TIT2L-GFP-ICL-TPT), characterized previously [7] was used for imaging experiments. Genotyping confirmed expression of Cre and tdTomato for this individual. The mouse was sacrificed at age P96 by trans-cardial perfusion with 4% paraformaldehyde. The brain was dissected and post-fixed in 4% paraformaldehyde at room temperature for 3-6 hr followed by overnight fixation at 4 deg. C. The brain was rinsed with 1× PBS and stored in 1× PBS with 0.1% sodium azide prior (Cat:S2002, Sigma-Aldrich) prior to sectioning. 200-µm thick cortical sections were cut on a vibratome and stored in 1× PBS. Prior to OTLS imaging, brain slices were incubated in a mixture of 68% 2,2′-thiodiethanol (TDE) (Cat:166782, Sigma-Aldrich) and 32% 1× PBS for clearing. The refractive index of the solution (*n* ~ 1.46) was verified using a refractometer (PA202, Misco). Procedures involving mice were approved by the Institutional Animal Care and Use Committee of the Allen Institute for Brain Science in accordance with NIH guidelines.

### Collection and processing of heart, lung, prostate, and lymph nodes

Lung, heart, prostate, and lymph nodes were collected from a CD11-YFP, Actin-dsRed expressing mouse. Tissues were fixed for 24 hr at 4 deg. C in 1 part fixative (Cat:554655, BD Biosciences) and 2 parts 1× PBS and incubated in blocking buffer for 24 hr at 37 deg. C. The buffer consisted of 30 mL Tris (Cat:252859, Sigma-Aldrich), 0.3 mL NMS (Cat:SML1128, Sigma-Aldrich), 0.3 mL BSA (Cat:A2058, Sigma-Aldrich), and 0.09 mL TritonX100 (Cat:T8787, Sigma-Aldrich). Lymph nodes were stained for 4 days at 37 deg. C in 400 μL blocking buffer, 2 μL CD3-BV421 (Cat: 100228, BioLegend), and 2 μL B220-e660 (Cat: 50-0452-82, ThermoFisher). Lung tissue was stained for 3 days at 37 deg. C in 500 μL blocking buffer and 2.5 μL Epcam-APC (Cat: 17-5791-82, ThermoFisher). Heart tissue was stained for 1 day with 1 mM DRAQ5. Prostate tissue was incubated with fluorophore-conjugated anti-CK8-18 (Cat:MS743S0, Fisher) conjugated to Alexa-Fluor 488 (Cat:A20181, Invitrogen) (1:100 dilution) in PBS/1% non-fat dry milk/0.2% Triton X-100 at 37 deg. C for 7 days with gentle agitation. All tissues were then cleared with the Ce3D solution, consisting of 14 mL of 40% N-methyl-acetamide (Cat:M26305, Sigma-Aldrich), 25 μL Triton X-100 (Cat:T8787, Sigma-Aldrich), 20 g Histodenz (Cat:D2158, Sigma-Aldrich), and 125 μL Thioglycerol (Cat:88640, Sigma-Aldrich) for 1 day at room temperature. Procedures involving mice were approved by the Institutional Animal Care and Use Committee of the University of Washington in accordance with NIH guidelines.

### Collection and processing of expanded mouse kidney

4% PFA fixed mouse kidney was sliced to 200 μm and processed using a previously described protocol [8]. The tissue was then incubated in blocking/permeabilization buffer for 6 hr at 4 deg. C. Primary antibodies goat anti-podocalyxin (cat: AF1556, R&D Sys. Inc., 1:50) and rabbit anti-collagen IV (cat: ab6586, abcam, 1:50) were diluted with blocking/permeabilization buffer and used to stain the tissue for 2 days at 4 deg. C. The tissue was then washed with 1× PBS three times at room temperature (1 hr each). Fluorescently-labeled secondary antibodies, Alexa 488 conjugated WGA (cat: W11261, Thermo Fisher Scientific, 1:25), and Hoechst 33342 were then diluted in blocking/permeabilization buffer to stain the tissue for 2 days at 4 deg. C. The tissue was washed with 1xPBS three times at room temperature (1 hr each) followed by incubating in 1 mM MA-NHS (cat:730300, Sigma-Aldrich) for 1 hr at room temperature. The tissue was then incubated in monomer solution for 1 hr at 4 deg. C and then gelled in a humidified environment at 37 deg. C for 2 hr. Excess gel was removed and the specimen was digested by proteinase K (cat: EO0491, Thermo Fisher Scientific) at 37 deg. C for two days and then collagenase (cat; C7926, Sigma-Aldrich) at 37 deg. C for two days refreshing the solution daily. After digestion, the specimen was incubated in DI water for at least 2 hr and the expansion factor was determined through measuring the dimensions of the gel. The expanded specimen was mounted on poly-lysine coated film for imaging.

### Collection and processing of human prostate biopsies

All specimens were obtained from an IRB-approved genitourinary biorepository with patient consent. Core-needle biopsy specimens were obtained from fresh *ex vivo* prostatectomy specimens using an 18-gauge (approximately 1 mm inner diameter) needle biopsy device (Bard Max Core, Bard Biopsy). The biopsy was immediately placed in 10% neutral buffered formalin, where it was maintained at room temperature for 24 hrs. In contrast to mouse tissues, we found that human tissues require more aggressive solvent-clearing approaches (**Supplementary Figure 21**). Due to its clearing efficacy and non-toxic nature, we used ECi-clearing, which we observed does not interfere with downstream histology or immunohistochemistry (**Supplementary Figure 22**).

Biopsies were then washed in 1× PBS with 0.1% Triton X-100 (Cat:T8787, Sigma-Aldrich), and each biopsy was stained for 4 hr in a 1:2000 dilution of TO-PRO3 Iodide (Cat:T3605, Thermo-Fischer) at room temperature with light shaking. Each biopsy was then dehydrated in ethanol for with 25/75, 50/50, 75/25, and 100/0 grades. The dehydration time for each grade was 1 hr, and the 100% ethanol grade was performed twice to ensure removal of any excess water. Biopsies were then stained in 1:2000 dilution of Eosin-Y (Cat:3801615, Leica Biosystems) for 4 hr at room temperature with light shaking. Finally, biopsies were cleared in ethyl-cinnamate (Cat:112372, Sigma-Aldrich) for 1 hr. Biopsy #12 was stained with anti-CK8. The biopsy issue was incubated simultaneously with fluorophore-conjugated anti-CK8-18 (Cat:MS743S0, Fisher) conjugated to Alexa-Fluor 488 (Cat:A20181, Invitrogen) (1:100 dilution) in PBS/1% non-fat dry milk/0.2% Triton X-100 at 37 deg. C for 7 days with gentle agitation.

## Supporting information

Summary Video

Supplementary ZEMAX Files

Supplementary CAD Files

## Data Availability

All raw and processed data generated in this work, including the images provided in the manuscript and supplementary material are available from the authors upon request. The customized CAD and ZEMAX files are available as **Supplementary CAD Files** and **Supplementary ZEMAX Files**.

## Code Availability

The computer code used to acquire, process, and generate the images in this study is available as **Supplementary Code Files**. **Supplementary Data** and instructions for installing and using the computer code are available at https://figshare.com/articles/Supplementary_Data/7685597.

## Acknowledgements

Human prostate specimens were provided by the GU Specimen Biorepository, University of Washington, which is supported by resources of the Department of Defense Prostate Cancer Research Program (W81XWH-14-2-0183), the Pacific Northwest Prostate Cancer SPORE (P50CA97186), a P01 NIH grant (PO1 CA163227), and the Institute for Prostate Cancer Research of the University of Washington. This work was funded in part by NIH grants R01 CA175391 (Liu), R01 DE023497 (Liu), F32 CA213615 (Glaser), the University of Washington Royalty Research Fund UW RRF A107248 (Liu), a UW CoMotion Innovation Award (Liu and Reder), a Safeway / NCI SPORE developmental award (subcontract) from the Fred Hutchinson Cancer Center (Liu and Dintzis), Prostate Cancer Foundation Young Investigator Award (Reder), the PCRP Idea Generation Award from Department of Defense, Congressionally Directed Medical Research Program (CDMRP) PC170176 (Liu and True), and the NIDDK Diabetic Complications Consortium grants DK076169 and DK115255 (Vaughan). Finally, the authors would like to thank the NVIDIA GPU Grant Program for donation of the GPUs used in this study.

## Contributions

A.G., N.R., P.N., M.G., J.V., L.T. and J.L. designed the studies. A.G. and J.L. designed the multi-immersion open-top light-sheet microscope. J.D. designed the multi-immersion collection objective and provided input on the microscope design. A.G., Y.C., C.Y, L.B., W.X., and L.W fabricated the microscope. A.G. and P.N. prepared and imaged the mouse brain slices. C.S., A.G., and E.M. prepared and imaged the mouse organs. A.H., C.M., and J.V. prepared the mouse kidney tissues. A.G., A.H., and C.M. imaged the expanded mouse kidney tissues. N.R., S.K., L.T., and E.M. prepared the human prostate tissues. A.G., S.K., and N.R. imaged the human prostate tissues. E.M. prepared all downstream histology for the study. N.R. and L.T. histologically characterized all human prostate tissues. All authors prepared the manuscript.

## Corresponding Author

Correspondence to <u>Adam K. Glaser</u> or <u>Jonathan T.C. Liu</u>

