## Supplementary Document for "Multi-immersion open-top light-sheet microscope for high-throughput imaging of cleared tissues"

**Supplementary Discussion**

*Advantages of open-top architecture over alternative designs*

Early light-sheet microscope architectures, known as selective plane illumination microscopy (SPIM), required single specimens to be carefully mounted in small capillary tubes and/or agarose, which were then suspended within a liquid-filled cuvette, a process that was well-suited for imaging small model organisms, but less ideal for imaging larger cleared tissues [9-11]. Subsequent LSFM architectures, including ultramicroscopy (UM), dual-inverted selective plane illumination microscopy (diSPIM), and light-sheet theta microscopy (LSTM), imaged specimens from above, with objectives dipped downward into a large liquid-filled chamber [12-14]. While these geometries are better suited for imaging large cleared tissues, submerging specimens in a liquid reservoir has several drawbacks. For example, the imaging objectives and specimen both share a common liquid reservoir, which must be replaced after each imaging session due to contamination by the specimen. This would also lead to inter-specimen contamination during multi-specimen drug screening studies. In addition, these systems are unable to image tissues cleared with certain highly corrosive organic solvents (e.g., dibenzyl-ether, benzyl-alcohol, and benzyl-benzoate) without the use of specialized immersion objectives. Finally, the cleared tissues equilibrate to a similar density as the surrounding liquid reservoir, making them prone to floating and therefore creating logistical challenges in terms of mounting and immobilizing specimens for large-volume imaging experiments (**Supplementary Figure 2**).

To address this need, open-top light-sheet (OTLS) microscopes have recently been developed [15-18]. The "open-top" mimics the geometry of a flatbed scanner and images the bottom side of specimens placed on a transparent surface. Since all optical components and hardware are located below the specimen, such a design can theoretically accommodate specimens of arbitrary size, shape, and numbers. The architecture is readily compatible with an array of specimen holders, which simplifies specimen mounting and facilitates high-throughput imaging. An added advantage of the open-top approach is that the specimen is physically separated from the imaging objectives by a transparent substrate/barrier, which overcomes several of the previously mentioned shortcomings and logistical difficulties of the UM, diSPIM, and LSTM systems.

While the OTLS architecture is more versatile, it also presents unique optical challenges, as the off-axis (45 deg.) illumination light sheet and collected fluorescence are not easily index-matched into and out of the specimen and holder, which can result in aberrations. To enable aberration-free imaging, both a water prism or solid-immersion lens (SIL) have been proposed to serve as an interface for the illumination and collection beams to transition from air into a higher-index medium: the former by providing a normal interface to mitigate off-axis aberrations, and the latter by providing a wavefront-matched interface to prevent both off-axis and spherical aberrations [15-17]. While these solutions address the issue of off-axis aberrations, they limit specimen scanning to 2D rather than a desired 3D (i.e., the specimens cannot be physically lowered without colliding with the water prism or SIL). In addition, these solutions are not readily compatible with a range of refractive indices, as the specimen must precisely match the refractive index of the water prism or SIL material. The multi-immersion open-top light-sheet microscope overcomes these limitations and improves upon our 1$^{st}$-generation OTLS prototype (**Supplementary Figure 3**).

*Refractive index considerations for various clearing protocols*

Precise refractive index matching of the immersion medium, holder, and sample is necessary for aberration-free imaging with the open-top architecture. This refractive index value is dictated by the clearing protocol used on the sample, and spans a wide range ($n$ = 1.33 – 1.56). These protocols can be categorized into aqueous, solvent, and expansion protocols [19-41].

Aqueous-based protocols involve optional lipid removal using detergents, followed by immersion in mixtures of a single or multiple water-soluble reagents. Because these mixtures contain some dilution of water, their exact refractive index value is difficult to control precisely (for the purposes of matching the index of the specimen holder material). For example, when initially clearing a specimen, the water content of the specimen will homogenize with the volume of refractive index matching media, potentially leading to a slightly different final refractive index than intended. Therefore, several successive refractive index matching immersion steps are necessary to achieve the intended specimen refractive index. In addition, evaporation of water content from the specimen can lead to a change in refractive index over time. To mitigate this, specimens should be covered during imaging.

Solvent-based protocols involve dehydration of tissue samples, and replace water content with organic reagents of a higher refractive index. Unlike aqueous protocols, the refractive index of solvent-cleared specimens is easier to control. However, similar to the issue of water content for aqueous-based protocols, the alcohol used for dehydration will homogenize with the final organic reagent. To achieve the desired final refractive index, we found that specimens should be cleared multiple times in the final organic reagent, or in a large enough volume of reagent such that the volume of alcohol within the specimen has a negligible effect on the specimen's final refractive index.

Finally, expansion-based protocols involve several steps which physically magnify samples by swelling a hydrogel. To date, these protocols have all used water ($n$ = 1.33) for the final swelling and expansion steps. Therefore, the refractive index of expanded specimens is easy to control. However, care must be taken to cover the gels during imaging and prevent them from drying out.

*Material selection and dispersion*

As mentioned in the main manuscript, diffraction-limited imaging performance is only achieved for optical path differences of $\Delta n \times t < 0.002$. However, the refractive index of materials and clearing reagents all vary with wavelength (i.e., dispersion) (see **Supplementary Figure 20**). The dispersion of glasses are well characterized and available from manufacturers and online resources. However, the dispersion of monomers and polymers are less characterized and more difficult to obtain. Finally, the dispersion of many clearing reagents has not been characterized. While more common reagents (e.g., water, TDE,

DMSO, and Glycerol) are available, the increasing complexity of clearing cocktails, particularly for aqueous protocols, makes it difficult to determine the final mixture's refractive index. For example, the latest CUBIC clearing protocol uses several reagents and antipyrine/N-methylnicotinamide, all of which not been characterized [42]. Therefore, future design and optimization of OTLS-based systems will require more precise characterization of the dispersion of various clearing reagents.

*Objective design for open-top imaging*

The performance of the OTLS system depends most critically on the choice of microscope objectives. For the current system, a multi-immersion objective designed by ASI / Special Optics was utilized. This particular objective has a specific numerical aperture (NA) and working distance (WD). The WD dictates the maximum imaging depth (*h*) of our system (see **Supplementary Figure 22**). However, future systems requiring either lower or higher resolution (with a resulting trade-off in field of view), and more or less imaging depth, could be designed around a different objective. For a given NA, the maximum imaging depth can be extended by increasing the WD of the objective, which usually increases the physical size of the objective and the diameter of the pupil near the back focal plane of the objective. The limiting factor to increasing the size of the objective would be cost and compatibility with other optical and mechanical components. The relationship between the NA and *h* is plotted in **Supplementary Figure 22**. Note that in practice, *h* will be less than the values shown since the mechanical housing of most objectives will protrude above the front element of the objective and field of view will reduce the usable working distance of the objective.

### Supplementary Table 1 - OTLS system components list

| Part Number | Vendor | Notes | Quantity |
|---|---|---|---|
| **Immersion Chamber** | | | |
| IMMERSION CHAMBER | Hilltop Technologies | CAD file provided | 1 |
| ILLUMINATION OBJECTIVE MOUNT | Hilltop Technologies | CAD file provided | 1 |
| COLLECTION OBJECTIVE MOUNT | Hilltop Technologies | CAD file provided | 1 |
| P8 | Thorlabs | Immersion chamber supports | 4 |
| PB4 | Thorlabs | Support post pedestals | 4 |
| **Illumination Objective** | | | |
| XLFLUOR 340/4X | Olympus | Illumination objective | 1 |
| LA4725-A | Thorlabs | Illumination SIL | 1 |
| 0.945 X 0.030 75 FLUORCARBON | Apple Rubber | SIL gasket | 1 |
| SM1RR | Thorlabs | Retaining ring for holding illumination SIL | 1 |
| **Collection Objective** | | | |
| MULTI-IMMERSION OBJECTIVE | ASI / Special Optics | Collection objective NA = 0.40 @ $n$ = 1.45 (range $n$ = 1.33 – 1.56) | 1 |
| 1.461 X 0.063 70 BUNA-N | Apple Rubber | Collection objective gasket | 1 |
| **Illumination Optics** | | | |
| AC127-019-A-ML | Thorlabs | Collimating lens $f$ = 19 mm | 1 |
| ACY254-50-A | Thorlabs | Cylindrical lens $f$ = 50 mm | 1 |
| ACY254-150-A | Thorlabs | Cylindrical lens $f$ = 150 mm | 1 |
| AC254-75-A-ML | Thorlabs | Relay lenses $f$ = 75 m | 2 |
| CLS-SL | Thorlabs | Scan lens $f$ = 70 mm | 1 |
| TTL200-A | Thorlabs | Tube lens $f$ = 200 mm | 1 |
| **Collection Optics** | | | |
| TTL165-A | Thorlabs | Tube lens $f$ = 165 mm | 1 |
| SM2F | Thorlabs | Adjustable collimation adapter for tube lens | 1 |
| **Imaging Camera** | | | |
| ORCA-FLASH4.0 V2 | Hamamatsu | sCMOS camera | 1 |
| **Filter Wheel** | | | |
| FW102C | Thorlabs | Motorized filter wheel | 1 |
| FF02-447/60-25 | Semrock | 405 bandpass filter | 1 |
| FF03-525/50-25 | Semrock | 488 bandpass filter | 1 |
| FF01-618/50-25 | Semrock | 561 bandpass filter | 1 |
| FF01-721/65-25 | Semrock | 638 bandpass filter | 1 |
| **Scanning Stage Components** | | | |
| MS-2000 XY STAGE | ASI | Motorized XY stage, modified top-plate for open-top mounting | 1 |
| LS-50 Z TRANSLATORS | ASI | Motorized Z axis translators | 2 |
| P4 | Thorlabs | Breadboard supports | 4 |
| PB4 | Thorlabs | Support post pedestals | 4 |

| | | | |
|---|---|---|---|
| TR2.5 | Thorlabs | LS50 raisers | 8 |
| PH2.5 | Thorlabs | LS50 raisers | 8 |
| **Customized Specimen Holders** | | | |
| FLEXVAT HOLDER AND STAGE ADAPTER | Flexvat.com | For holding FEP film and expanded samples (*n* = 1.33) CAD file provided | 1 |
| FLAT PLATE STAGE ADAPTER | Hilltop Technologies | For holding fused silica plate (*n* = 1.46) CAD file provided | 1 |
| WELL PLATE STAGE ADAPTER | Hilltop Technologies | For holding well plate with PMMA bottom (*n* = 1.49) CAD file provided | 1 |
| BIOPSY STAGE ADAPTER | Hilltop Technologies | For holding HIVEX biopsy holder (*n* = 1.56) CAD file provided | 1 |
| HIVEX HOLDER | In house | For holding biopsies (*n* = 1.56) CAD file provided | 1 |
| **Laser Package** | | | |
| 90420 | Cobolt | Skyra fiber-coupled 405, 488, 561, 638 lasers | 1 |
| 12422 | Cobolt | Heatsink | 1 |
| PM-S405-XP-CUSTOM | Thorlabs | Custom FC/APC FC/PC S405-XP fiber | 1 |
| **Scanning Mirror** | | | |
| 6210H | Cambridge Technologies | Galvanometer mirror | 1 |
| 6210H Mount | Cambridge Technologies | Mirror mount | 1 |
| NI-6115 | National Instruments | Waveform generator | 1 |
| **Optomechanical Assembly Components** | | | |
| SM1A61 | Thorlabs | - | 1 |
| SM1ZM | Thorlabs | - | 2 |
| SM1A24 | Thorlabs | - | 1 |
| SM1L05 | Thorlabs | - | 1 |
| SM1Z | Thorlabs | - | 1 |
| CXY1 | Thorlabs | - | 1 |
| LCP02 | Thorlabs | - | 8 |
| CRM1-P | Thorlabs | - | 2 |
| CPB1 | Thorlabs | - | 5 |
| LCP01 | Thorlabs | - | 9 |
| CP02 | Thorlabs | - | 1 |
| KCB2EC | Thorlabs | - | 4 |
| PFE20-P01 | Thorlabs | - | 5 |
| ER3 | Thorlabs | - | 5 |
| ER12 | Thorlabs | - | 8 |
| ER10 | Thorlabs | - | 4 |
| ER4 | Thorlabs | - | 16 |
| ER2 | Thorlabs | - | 8 |
| SM1FC | Thorlabs | - | 1 |
| SM1A6 | Thorlabs | - | 1 |

| | | | |
|---|---|---|---|
| C6W | Thorlabs | - | 1 |
| LB4C | Thorlabs | - | 1 |
| SM2F | Thorlabs | - | 1 |
| SM2A55 | Thorlabs | - | 1 |
| TR075 | Thorlabs | - | 3 |
| PH1 | Thorlabs | - | 3 |
| MF469-35 | Thorlabs | - | 1 |
| PF175 | Thorlabs | - | 6 |
| AP45 | Thorlabs | - | 8 |
| SM1A12 | Thorlabs | - | 1 |
| M32M34S | Thorlabs | - | 1 |
| LCPB1 | Thorlabs | - | 4 |
| SM2A20 | Thorlabs | - | 1 |
| KCB1E | Thorlabs | - | 1 |
| SMA2A55 | Thorlabs | - | 1 |
| MB1236 | Thorlabs | - | 2 |
| TR2 | Thorlabs | - | 1 |
| PH1.5 | Thorlabs | - | 1 |

### Supplementary Table 2 - Computer hardware specifications

| Part Number | Vendor | Notes | Quantity |
|---|---|---|---|
| **Acquisition Computer** | | | |
| Precision Tower 5810 | Dell | 1x CPU<br>2 PCI-E 3.0 x16 (double-width) slots,<br>1 PCI-E 3.0 x8 (single-width) slots,<br>1 PCI-E 2.0 x4 slot<br>1 PCI-E 2.0 x1 slot<br>4 3.5" drive bays<br>USB3.0 (1 front port)<br>USB2.0 (3 front ports)<br>USB3.0 (3 rear ports)<br>USB2.0 (3 internal ports) | 1 |
| SSD 2.00 TB 960 PRO Series | Samsung | 2.5" SATA 6.0Gb/s Solid State Drive (RAID0) | 4 |
| MegaRAID 9361-8i | LSI | SAS 12Gb/s PCIe 3.0 8-Port Controller with 1GB Cache (1x internal RAID0 arrays) | 1 |
| GPU Quadro K620 | NVIDIA | 12 GB GDDR5X (administrator account) | 1 |
| 10-Gigabit Ethernet Adapter MCX311A | Mellanox | ConnectX-3 EN MCX311A (1x SFP+) | 1 |
| RAM KVR21E15D8/16 | Kingston | 16 GB ECC Registered DDR4 2133 PC4 1700 | 2 |
| FireBird 1xCLD-2PE8 | Active Silicon | PCIe 3.0 x8 sCMOS camera frame grabber | 1 |
| Windows 7 | Microsoft | 64-bit | 1 |
| **Computing Server** | | | |
| SuperWorkstation 7049GP-TRT | SuperMicro | 2x CPUs<br>4 PCI-E 3.0 x16 (double-width) slots,<br>2 PCI-E 3.0 x16 (single-width) slots,<br>1 PCI-E 3.0 x4 (in x8) slot<br>8 Hot-swap 3.5" drive bays<br>Up to 2TB ECC 3DS LRDIMM, up to DDR4-2666MHz; 16 DIMM slots<br>Dual socket P (LGA 3647) supports Intel® Xeon® Scalable Processors | 1 |
| CPU Xeon Gold 6134 | Intel | 8-core 3.20GHz 24.75MB Cache (130W) | 2 |
| NVMe SSD 960 PRO M.2 | Samsung | 512GB PCIe 3.0 x4 NVMe (1 Cache, 1 OS) | 2 |
| SSD 1.92TB 5200 ECO Series | Micron | 2.5" SATA 6.0Gb/s Solid State Drive (RAID0) | 4 |
| SSD 2.00 TB 960 PRO Series | Samsung | 2.5" SATA 6.0Gb/s Solid State Drive (RAID0) | 4 |
| MegaRAID 9361-8i | LSI | SAS 12Gb/s PCIe 3.0 8-Port Controller with 1GB Cache (2x internal RAID0 arrays) | 1 |
| MegaRAID 9380-8e | LSI | SAS 12Gb/s PCIe 3.0 8-Port Controller with 1GB Cache (1x external RAID6 array) | 1 |
| 10-Gigabit Ethernet Adapter MCX311A | Mellanox | ConnectX-3 EN MCX311A (1x SFP+) | 1 |
| RAM KTD-PE421LQ/32G | Kingston | 32 GB ECC Registered DDR4 2133 PC4 1700 | 12 |
| GPU TitanXP | NVIDIA | 12 GB GDDR5X (administrator account) | 1 |
| GPU Quadro P6000 | NVIDIA | 24 GB GDDR5X (discrete device assignment guest account) | 1 |
| OS Windows Server 2016 | Microsoft | 64-bit | 1 |

| Direct-attached Storage | | | |
|---|---|---|---|
| Chasis STX-3316 3U | Thinkmate | 16x Hot-Swap 3.5" SATA/SAS3<br>12Gb/s SAS Single Expander | 1 |
| Exos 7E8 Series (512e) | Seagate | 8.0TB SAS 3.0 12.0Gb/s 7200RPM - 3.5" (RAID6 array 96 TB total storage) | 16 |
| External SAS Cable | Thinkmate | 1-meter 12Gb/s to 12Gb/s SAS - SFF-8644 to SFF-8644 | 1 |

### Supplementary Table 3 - Imaging datasets

| Dataset Name | Sampling (µm/px) | Dimensions (px) | Size (mm) | Fluorescent Markers | Laser (mW) | Attenuation $(mm^{-1})$ | Time (min) | Size (GB) | B3D (GB) |
|---|---|---|---|---|---|---|---|---|---|
| **TDE clearing (*n* = 1.46)** | | | | | | | | | |
| Brain Slice (1) | 0.342 (X)<br>0.485 (Y)<br>0.342 (Z) | 14912<br>17920<br>624 | 4.9 (X)<br>8.7 (Y)<br>0.2 (Z) | GFP (488) | 6 mW | 1.0 | 13 | 172 | 24 |
| Brain Slice (2) | 0.342 (X)<br>0.485 (Y)<br>0.342 (Z) | 14912<br>17920<br>624 | 4.9 (X)<br>8.7 (Y)<br>0.2 (Z) | GFP (488) | 6 mW | 1.0 | 13 | 172 | 25 |
| Brain Slice (3) | 0.342 (X)<br>0.485 (Y)<br>0.342 (Z) | 14912<br>17920<br>624 | 4.9 (X)<br>8.7 (Y)<br>0.2 (Z) | GFP (488) | 6 mW | 1.0 | 13 | 172 | 22 |
| Brain Slice (4) | 0.342 (X)<br>0.485 (Y)<br>0.342 (Z) | 14912<br>17920<br>624 | 4.9 (X)<br>8.7 (Y)<br>0.2 (Z) | GFP (488) | 6 mW | 1.0 | 13 | 172 | 22 |
| Brain Slice (5) | 0.342 (X)<br>0.485 (Y)<br>0.342 (Z) | 14912<br>17920<br>624 | 4.9 (X)<br>8.7 (Y)<br>0.2 (Z) | GFP (488) | 6 mW | 1.0 | 13 | 172 | 22 |
| Brain Slice (6) | 0.342 (X)<br>0.485 (Y)<br>0.342 (Z) | 14912<br>17920<br>624 | 4.9 (X)<br>8.7 (Y)<br>0.2 (Z) | GFP (488) | 6 mW | 1.0 | 13 | 172 | 23 |
| Brain Slice (7) | 0.342 (X)<br>0.485 (Y)<br>0.342 (Z) | 14912<br>17920<br>624 | 4.9 (X)<br>8.7 (Y)<br>0.2 (Z) | GFP (488) | 6 mW | 1.0 | 13 | 172 | 26 |
| Brain Slice (8) | 0.342 (X)<br>0.485 (Y)<br>0.342 (Z) | 14912<br>17920<br>624 | 4.9 (X)<br>8.7 (Y)<br>0.2 (Z) | GFP (488) | 6 mW | 1.0 | 13 | 172 | 22 |
| **Ce3D clearing (*n* = 1.49)** | | | | | | | | | |
| Mouse Lung | 0.335 (X)<br>0.471 (Y)<br>0.335 (Z) | 12261 (X)<br>24030 (Y)<br>2686 (Z) | 4.1 (X)<br>11.3 (Y)<br>0.9 (Z) | F-actin (561)<br>EpCAM (638) | 10 mW<br>1 mW | 1.0<br>1.0 | 118 | 3165 | 473 |
| Mouse Heart | 0.670 (X)<br>0.942 (Y)<br>0.670 (Z)<br>(2x binning) | 6617 (X)<br>6744 (Y)<br>5970 (Z) | 4.4 (X)<br>6.4 (Y)<br>4.0 (Z) | AF (488)<br>DRAQ5 (638) | 20 mW<br>2 mW | 1.0<br>1.0 | 41 | 532 | 67 |
| Mouse Prostate | 0.335 (X)<br>0.471 (Y)<br>0.335 (Z) | 14464 (X)<br>10688 (Y)<br>1472 (Z) | 4.8 (X)<br>5.0 (Y)<br>0.5 (Z) | CK8 (638) | 8 mW | 1.0 | 18 | 228 | 32 |
| Lymph Node (1) | 0.335 (X)<br>0.471 (Y)<br>0.335 (Z) | 5248 (X)<br>2880 (Y)<br>1472 (Z) | 1.8 (X)<br>1.4 (Y)<br>0.5 (Z) | CD3 (405)<br>CD11 (488)<br>F-actin (561)<br>B220 (638) | 2 mW<br>8 mW<br>1 mW<br>20 mW | 1.0<br>1.0<br>1.0<br>1.0 | 9 | 89 | 14 |
| Lymph Node (2) | 0.335 (X)<br>0.471 (Y)<br>0.335 (Z) | 6272 (X)<br>2880 (Y)<br>1472 (Z) | 2.1 (X)<br>1.4 (Y)<br>0.5 (Z) | CD3 (405)<br>CD11 (488)<br>F-actin (561)<br>B220 (638) | 2 mW<br>8 mW<br>1 mW<br>20 mW | 1.0<br>1.0<br>1.0<br>1.0 | 10 | 106 | 15 |
| Lymph Node (3) | 0.335 (X)<br>0.471 (Y)<br>0.335 (Z) | 6272 (X)<br>2944 (Y)<br>1472 (Z) | 2.1 (X)<br>1.4 (Y)<br>0.5 (Z) | CD3 (405)<br>CD11 (488)<br>F-actin (561)<br>B220 (638) | 2 mW<br>8 mW<br>1 mW<br>20 mW | 1.0<br>1.0<br>1.0<br>1.0 | 10 | 110 | 16.7 |

| | | | | | | | | | |
|---|---|---|---|---|---|---|---|---|---|
| Lymph Node (4) | 0.335 (X)<br>0.471 (Y)<br>0.335 (Z) | 5248 (X)<br>2880 (Y)<br>1984 (Z) | 1.8 (X)<br>1.4 (Y)<br>0.7 (Z) | CD3 (405)<br>CD11 (488)<br>F-actin (561)<br>B220 (638) | 2 mW<br>8 mW<br>1 mW<br>20 mW | 1.0<br>1.0<br>1.0<br>1.0 | 12 | 120 | 18 |
| Lymph Node (5) | 0.335 (X)<br>0.471 (Y)<br>0.335 (Z) | 6656 (X)<br>2880 (Y)<br>2432 (Z) | 2.2 (X)<br>1.5 (Y)<br>0.8 (Z) | CD3 (405)<br>CD11 (488)<br>F-actin (561)<br>B220 (638) | 2 mW<br>8 mW<br>1 mW<br>20 mW | 1.0<br>1.0<br>1.0<br>1.0 | 17 | 186 | 24 |
| Lymph Node (6) | 0.335 (X)<br>0.471 (Y)<br>0.335 (Z) | 6144 (X)<br>2944 (Y)<br>1984 (Z) | 2.1 (X)<br>1.5 (Y)<br>0.7 (Z) | CD3 (405)<br>CD11 (488)<br>F-actin (561)<br>B220 (638) | 2 mW<br>8 mW<br>1 mW<br>20 mW | 1.0<br>1.0<br>1.0<br>1.0 | 14 | 144 | 19 |
| Lymph Node (7) | 0.335 (X)<br>0.471 (Y)<br>0.335 (Z) | 4352 (X)<br>2880 (Y)<br>1984 (Z) | 1.5 (X)<br>1.5 (Y)<br>0.7 (Z) | CD3 (405)<br>CD11 (488)<br>F-actin (561)<br>B220 (638) | 2 mW<br>8 mW<br>1 mW<br>20 mW | 1.0<br>1.0<br>1.0<br>1.0 | 11 | 99 | 14 |
| Lymph Node (8) | 0.335 (X)<br>0.471 (Y)<br>0.335 (Z) | 4864 (X)<br>2880 (Y)<br>1920 (Z) | 1.6 (X)<br>1.5 (Y)<br>0.7 (Z) | CD3 (405)<br>CD11 (488)<br>F-actin (561)<br>B220 (638) | 2 mW<br>8 mW<br>1 mW<br>20 mW | 1.0<br>1.0<br>1.0<br>1.0 | 11 | 70 | 9 |
| **ECi clearing (*n* = 1.56)** | | | | | | | | | |
| Prostate Biopsy (1) | 0.321 (X)<br>0.454 (Y)<br>0.321 (Z) | 56872 (X)<br>2064 (Y)<br>2049 (Z) | 18.2 (X)<br>0.9 (Y)<br>0.6 (Z) | Eosin (561)<br>TO-PRO3 (638) | 1 mW<br>1 mW | 0.7<br>0.7 | 25 | 962 | 122 |
| Prostate Biopsy (2) | 0.321 (X)<br>0.454 (Y)<br>0.321 (Z) | 50155 (X)<br>2063 (Y)<br>2049 (Z) | 16.1 (X)<br>0.9 (Y)<br>0.6 (Z) | Eosin (561)<br>TO-PRO3 (638) | 1 mW<br>1 mW | 0.7<br>0.7 | 23 | 848 | 108 |
| Prostate Biopsy (3) | 0.321 (X)<br>0.454 (Y)<br>0.321 (Z) | 39564 (X)<br>2074 (Y)<br>2048 (Z) | 12.7 (X)<br>0.9 (Y)<br>0.6 (Z) | Eosin (561)<br>TO-PRO3 (638) | 1 mW<br>1 mW | 0.7<br>0.7 | 18 | 669 | 90 |
| Prostate Biopsy (4) | 0.321 (X)<br>0.454 (Y)<br>0.321 (Z) | 47364 (X)<br>2066 (Y)<br>2039 (Z) | 15.2 (X)<br>0.9 (Y)<br>0.6 (Z) | Eosin (561)<br>TO-PRO3 (638) | 1 mW<br>1 mW | 0.7<br>0.7 | 22 | 801 | 121 |
| Prostate Biopsy (5) | 0.321 (X)<br>0.454 (Y)<br>0.321 (Z) | 53271 (X)<br>2061 (Y)<br>2047 (Z) | 17.1 (X)<br>0.9 (Y)<br>0.6 (Z) | Eosin (561)<br>TO-PRO3 (638) | 1 mW<br>1 mW | 0.7<br>0.7 | 24 | 901 | 130 |
| Prostate Biopsy (6) | 0.321 (X)<br>0.454 (Y)<br>0.321 (Z) | 44860 (X)<br>2066 (Y)<br>2040 (Z) | 14.4 (X)<br>0.9 (Y)<br>0.6 (Z) | Eosin (561)<br>TO-PRO3 (638) | 1 mW<br>1 mW | 0.7<br>0.7 | 21 | 758 | 104 |
| Prostate Biopsy (7) | 0.321 (X)<br>0.454 (Y)<br>0.321 (Z) | 49844 (X)<br>2062 (Y)<br>2040 (Z) | 16.0 (X)<br>0.9 (Y)<br>0.6 (Z) | Eosin (561)<br>TO-PRO3 (638) | 1 mW<br>1 mW | 0.7<br>0.7 | 23 | 843 | 106 |
| Prostate Biopsy (8) | 0.321 (X)<br>0.454 (Y)<br>0.321 (Z) | 51714 (X)<br>2066 (Y)<br>2048 (Z) | 16.6 (X)<br>0.9 (Y)<br>0.6 (Z) | Eosin (561)<br>TO-PRO3 (638) | 1 mW<br>1 mW | 0.7<br>0.7 | 24 | 874 | 110 |
| Prostate Biopsy (9) | 0.321 (X)<br>0.454 (Y)<br>0.321 (Z) | 49221 (X)<br>2069 (Y)<br>2040 (Z) | 15.8 (X)<br>0.9 (Y)<br>0.6 (Z) | Eosin (561)<br>TO-PRO3 (638) | 1 mW<br>1 mW | 0.7<br>0.7 | 22 | 832 | 124 |
| Prostate Biopsy (10) | 0.321 (X)<br>0.454 (Y)<br>0.321 (Z) | 56075 (X)<br>2062 (Y)<br>2042 (Z) | 18.0 (X)<br>0.9 (Y)<br>0.6 (Z) | Eosin (561)<br>TO-PRO3 (638) | 1 mW<br>1 mW | 0.7<br>0.7 | 25 | 948 | 119 |

| | | | | | | | | | |
|---|---|---|---|---|---|---|---|---|---|
| Prostate Biopsy (11) | 0.321 (X)<br>0.454 (Y)<br>0.321 (Z) | 45483 (X)<br>2066 (Y)<br>2046(Z) | 14.6 (X)<br>0.9 (Y)<br>0.6 (Z) | Eosin (561)<br>TO-PRO3 (638) | 1 mW<br>1 mW | 0.7<br>0.7 | 22 | 769 | 99 |
| Prostate Biopsy (12) | 0.321 (X)<br>0.454 (Y)<br>0.321 (Z) | 47975 (X)<br>2060 (Y)<br>2040 (Z) | 15.4 (X)<br>0.9 (Y)<br>0.6 (Z) | CK8 (488)<br>TO-PRO3 (638) | 10 mW<br>1 mW | 0.7<br>0.7 | 23 | 811 | 102 |
| **ExM (*n* = 1.33)** | | | | | | | | | |
| Mouse Kidney (1) | 1.51 (X)<br>2.13 (Y)<br>1.51 (Z)<br>(4x binning) | 21192 (X)<br>9859 (Y)<br>728 (Z) | 32.0 (Y)<br>21.0 (X)<br>1.1 (Z) | DAPI (405)<br>WGA-lectin (488)<br>Podxl (561)<br>Coll IV (638) | 1 mW<br>30 mW<br>50 mW<br>50 mW | 2.0<br>2.0<br>2.0<br>2.0 | 64 | 608 | 84 |
| Mouse Kidney (2) | 0.377 (X)<br>0.533 (Y)<br>0.377 (Z) | 4884 (X)<br>4018 (Y)<br>2918 (Z) | 1.8 (X)<br>2.1 (Y)<br>1.1 (Z) | DAPI (405)<br>WGA-lectin (488)<br>Podxl (561)<br>Coll IV (638) | 1 mW<br>30 mW<br>50 mW<br>50 mW | 2.0<br>2.0<br>2.0<br>2.0 | 26 | 229 | 30 |

### Supplementary Figures

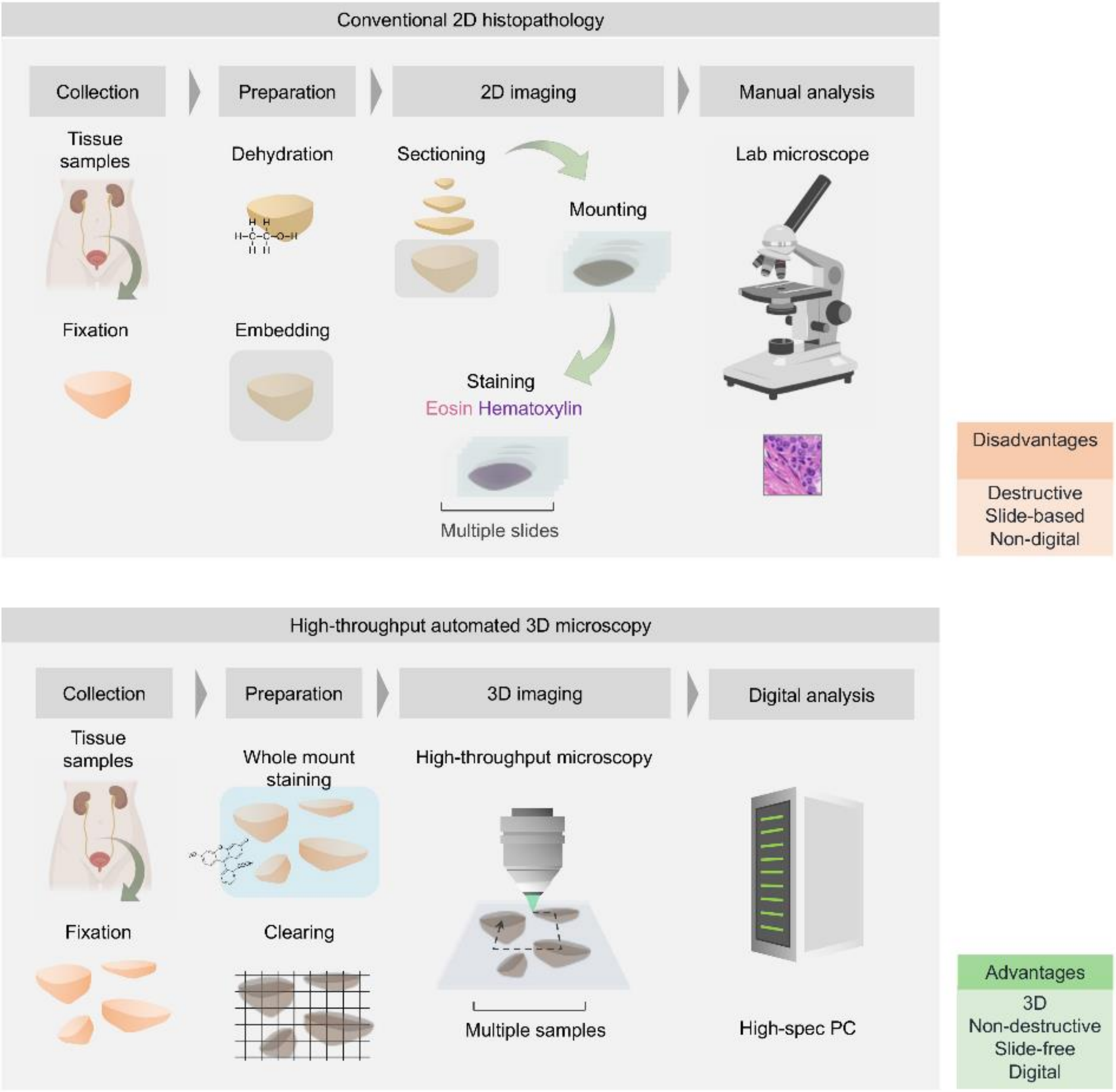


**Supplementary Figure 1. Comparison of 2D histopathology and 3D microscopy.** The conventional histopathology protocol requires specimens to be fixed, dehydrated, embedded in paraffin wax, physically sectioned into thin slices, and stained with hematoxylin and eosin (H&E). Slides are then manually analyzed with a conventional laboratory microscope. With the multi-immersion OTLS system, multiple specimens are collected, fixed, stained with fluorescent dyes, and cleared (using an aqueous, solvent, or expansion protocol). Multiple samples are then mounted on the system and imaged in an automated manner. The resulting 3D data is then digitally analyzed using a computer. In comparison to conventional histopathology, this workflow is 3D, non-destructive, slide-free, and inherently digital. Graphic adapted from [43]

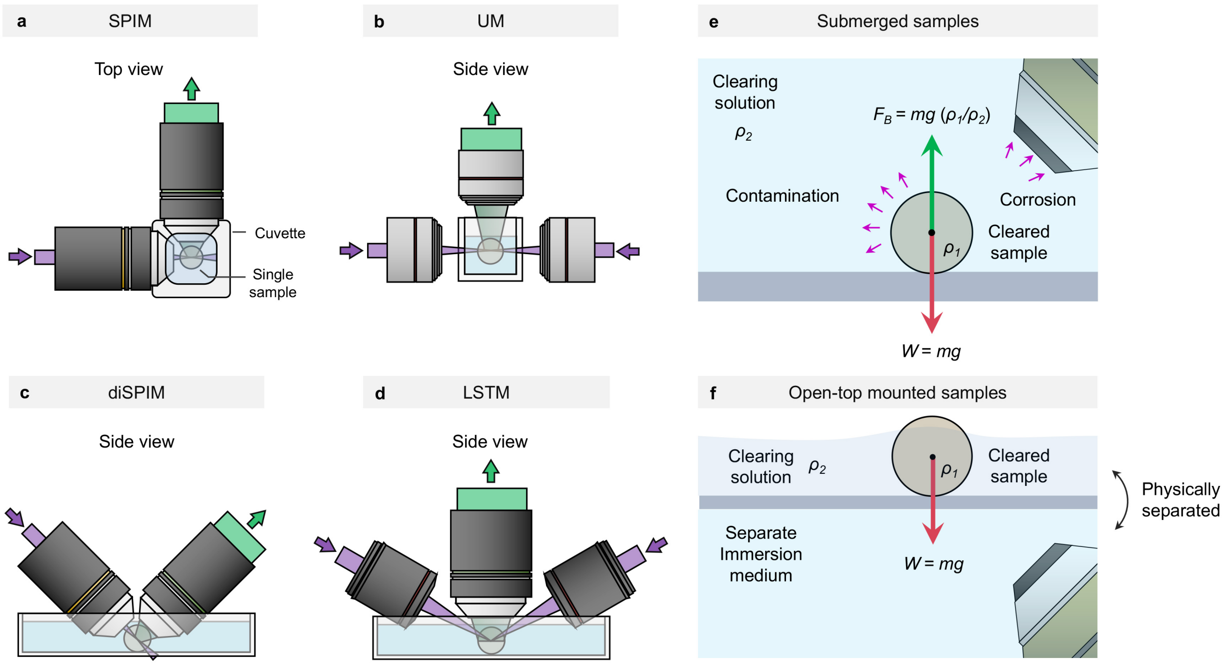


**Supplementary Figure 2. Alternative LSFM architectures. (a-d)** The geometry of SPIM, UM, diSPIM, and LSTM systems are shown. SPIM and UM systems place constraints on specimen size, shape, and number, limiting the ability to do automated high-throughput imaging. **(e)** diSPIM and LSTM systems require samples be submerged in a large liquid reservoir. In comparison to open-top mounted samples, this has several disadvantages **(f)**. The imaging objectives and sample both share a common liquid reservoir, which must be replaced after each imaging session due to contamination by the sample. In addition, these systems are unable to image tissues cleared with certain highly corrosive organic solvents (e.g., dibenzyl-ether, benzyl-alcohol, and benzyl-benzoate) without the use of specialized immersion objectives. Finally, the cleared tissues equilibrate to a similar density as the surrounding liquid reservoir, making them prone to floating and therefore creating logistical challenges in terms of mounting and immobilizing samples for large-volume imaging experiments. See **Supplementary Discussion**.

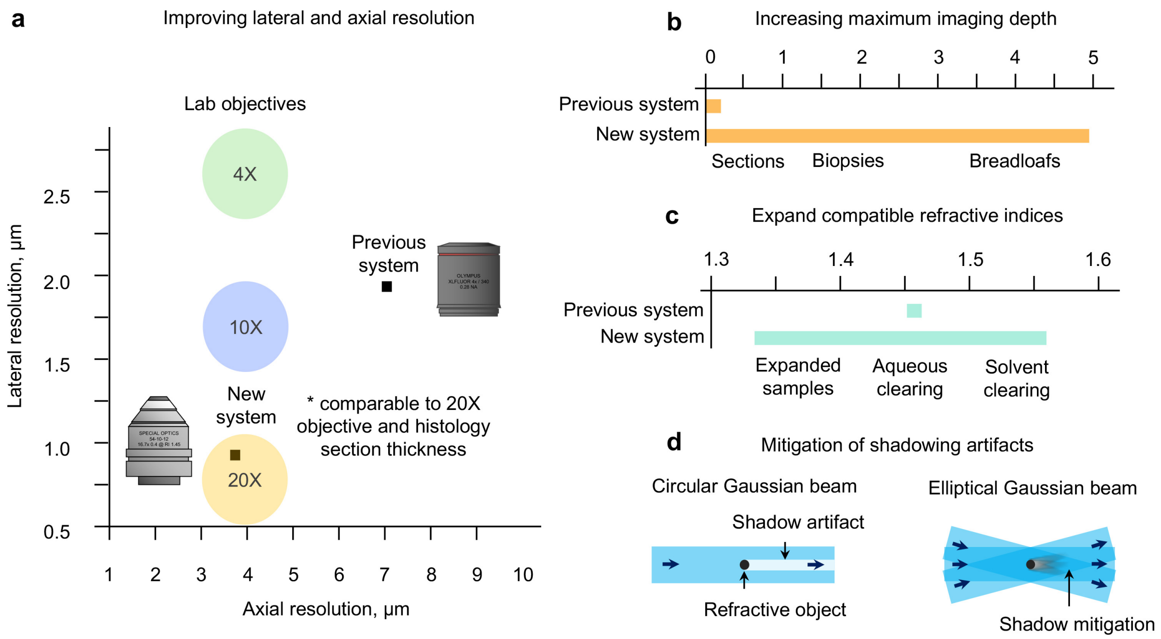


**Supplementary Figure 3. System improvements.** In comparison to a previous prototype system [16], our new OTLS system exhibits **(a)** improved axial and lateral resolution (an order-of-magnitude smaller focal volume), **(b)** ~20X greater imaging depth, **(c)** multi-immersion capabilities (n = 1.33 – 1.56), and **(d)** shadow mitigation.

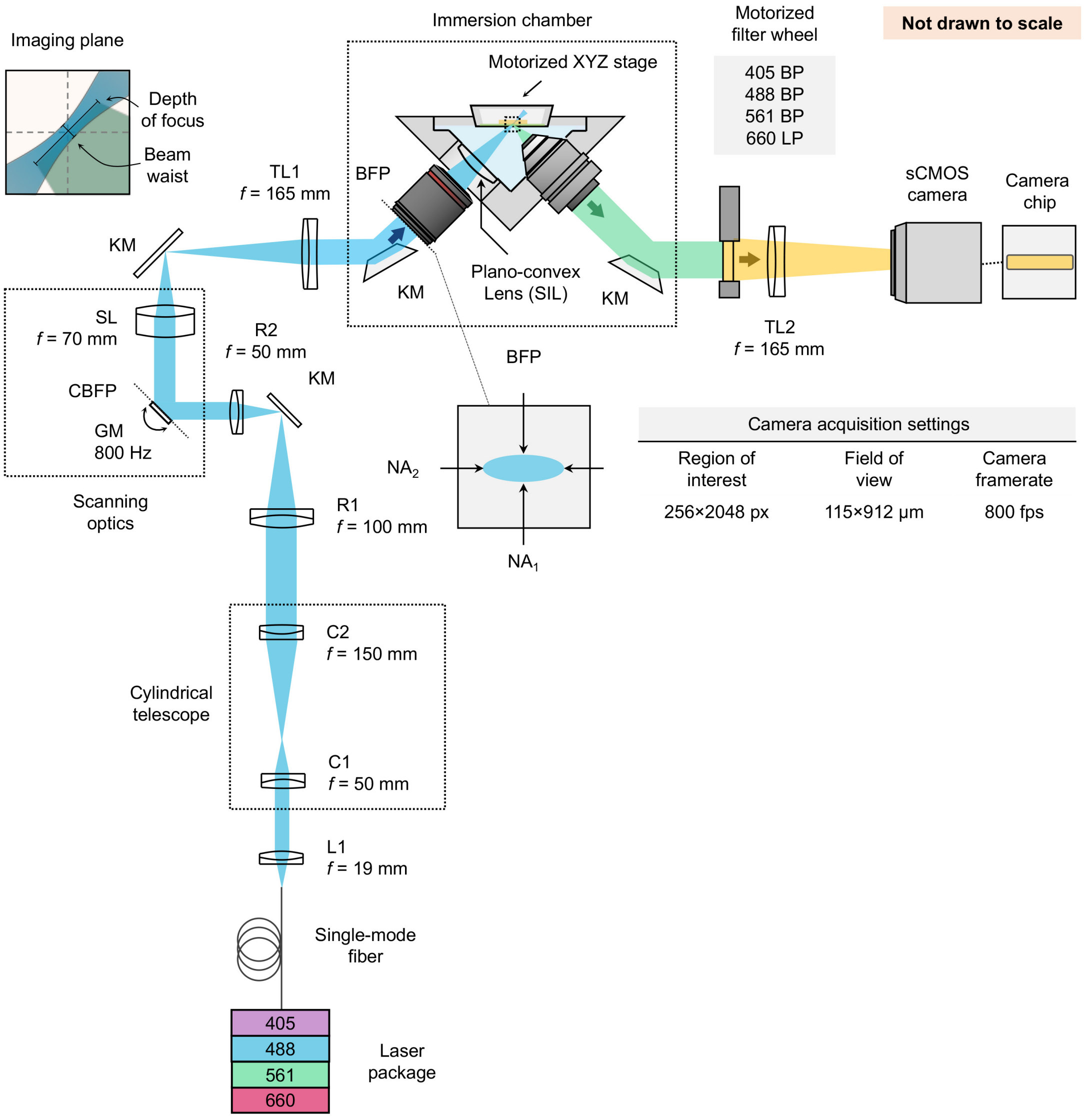


**Supplementary Figure 4. Optical layout.** Illumination light is coupled into the system by a single-mode fiber with a numerical aperture of 0.12 from a four-channel laser package, collimated with a lens, L1 ($f$ = 19 mm), and then expanded along one axis using a 3× cylindrical telescope consisting of lenses, C1($f$ = 50 mm) and C2 ($f$ = 150 mm) to provide multi-directional illumination. The resulting elliptical Gaussian beam is then relayed to the scanning galvanometer, GM (6210H, Cambridge Technology) using lenses R1 ($f$ = 100 mm) and R2 ($f$ = 50 mm). The scanning mirror is driven by a sinusoidal voltage from a waveform generator (PCI-6115, National Instruments) at a frequency of 800 Hz. The scanning beam is relayed to the back focal plan of the illumination objective (XLFLUOR340/4×, Olympus) using a scan lens, SL ($f$ = 70 mm) and tube lens, TL1 ($f$ = 165 mm). Finally, the elliptical beam travels through a plano-convex lens ($R$ = 34.5 mm), immersion media, holder, and finally sample. Fluorescence is collected by a multi-immersion objective (#54-10-12, Applied Scientific Instrumentation - ASI) which provides <1 µm in-plane resolution for all immersion media and filtered with a motorized filter wheel with band-pass filters for the 405 nm, 488 nm, 561 nm, and 638 nm excitation wavelengths. The filtered fluorescence is focused onto a 2048×2048-pixel sCMOS camera by a tube lens, TL2 ($f$ = 165 mm). The tube lens provides a Nyquist sampling of ~0.45 µm/pixel, which provides a horizontal field of view of ~0.9 mm over the 2048 pixels of the camera. The vertical field of view is reduced to 256 pixels match the depth of focus of the illumination light sheet (~110 µm). The 256 pixels are oriented parallel to the rolling shutter readout direction of the camera, which provides an exposure time of 1.25 ms and a framerate of 800 Hz.

Illumination optics ZEMAX model

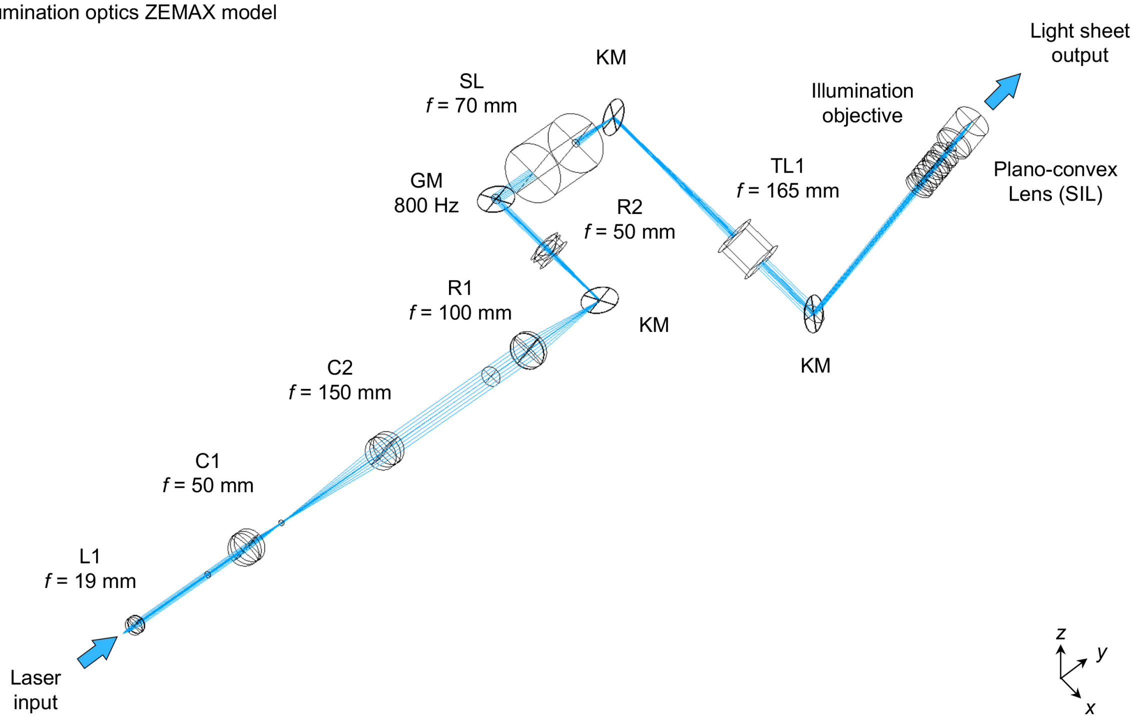


**Supplementary Figure 5. Illumination optics model.** 3D view of the illumination model, provided in **Supplementary ZEMAX files**. The illumination objective was modeled based on a patent [44]. All other components were modeled using manufacturer provided ZEMAX lens and blackbox files.

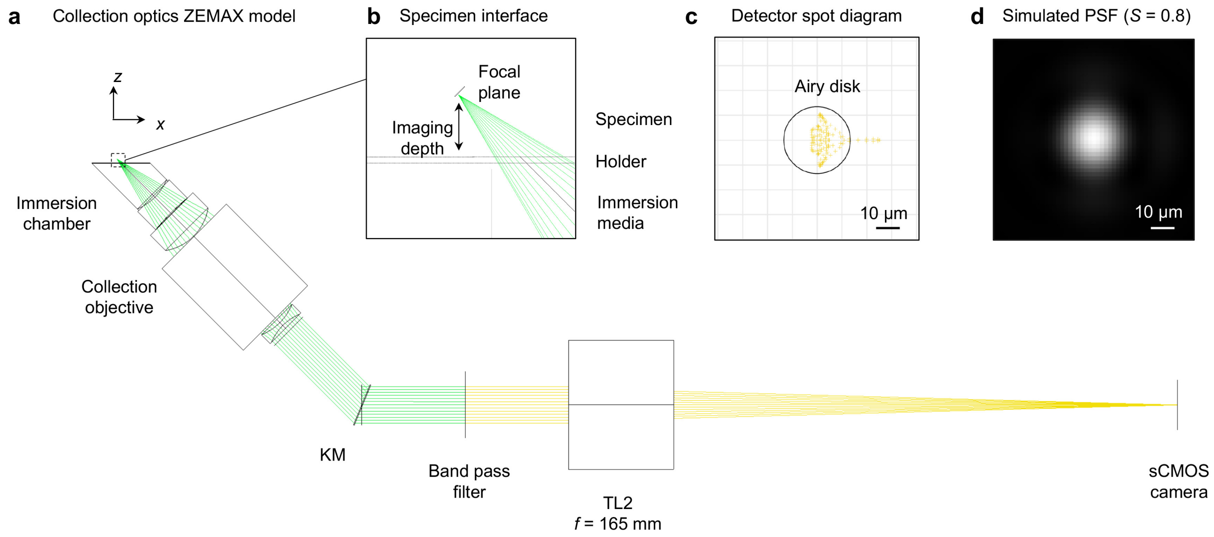


**Supplementary Figure 6. Collection optics model. (a)** ZEMAX model of the collection optics, provided in **Supplementary ZEMAX files**. A zoom-in of the specimen interface is shown in **(b)**. A representative detector spot diagram (ray-tracing) and simulated PSF (diffraction theory) are shown in **(c)** and **(d)**. This model file was used to calculate the Strehl Ratio (*S*) as a function of optical path difference plot in **Figure 2** of the main manuscript.

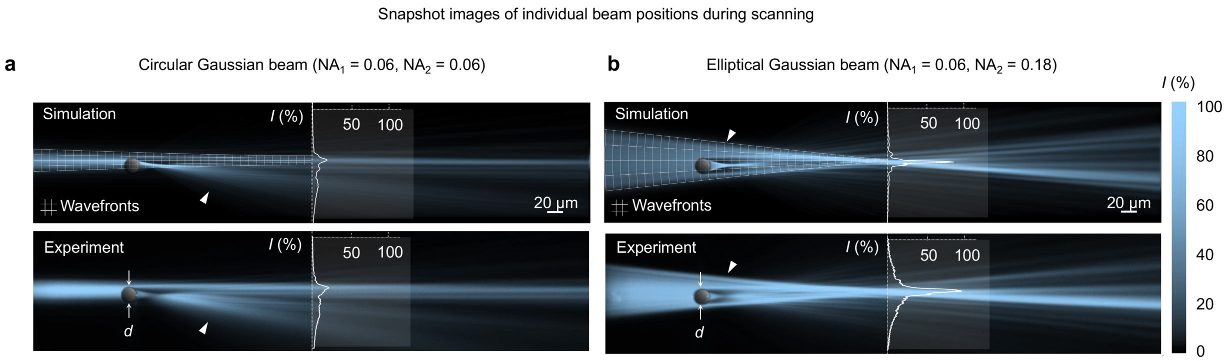


**Supplementary Figure 7. Multidirectional illumination. (a)** In comparison to conventional DSLM systems which utilize circular Gaussian beams, our OTLS system uses the mDSLM architecture and an elliptical Gaussian beam for mitigation of shadowing artifacts [1]. **(b)** Simulation and corresponding experimental image of a circular Gaussian beam ($NA_1 = NA_2 = 0.06$) propagating around a large glass sphere (diameter $d$ = 20 μm, $n_{sphere}$ = 1.59) embedded within a fluorescent gel ($n_{gel}$ = 1.46). The sphere is positioned at a depth of $z_{sphere}$ = 125 μm at an offset of $\Delta y$ = 2 μm from the optical axis of the pencil beam. The pencil beam focus is located at a depth of $z_{focus}$ = 350 μm. For a circular Gaussian beam (DSLM), the intensity at the beam focus is reduced by >75% relative to an unobstructed beam, as illustrated by the overlaid line profiles. **(c)** Simulation and corresponding experimental image of an elliptical Gaussian beam ($NA_1$ = 0.06, $NA_2$ = 0.18) propagating through an identical fluorescent gel and glass sphere. In contrast to the circular Gaussian beam used in DSLM, the increased angular diversity in the $y$ direction enables the elliptical Gaussian beam to experience only a ~10% reduction in intensity relative to an unobstructed beam.

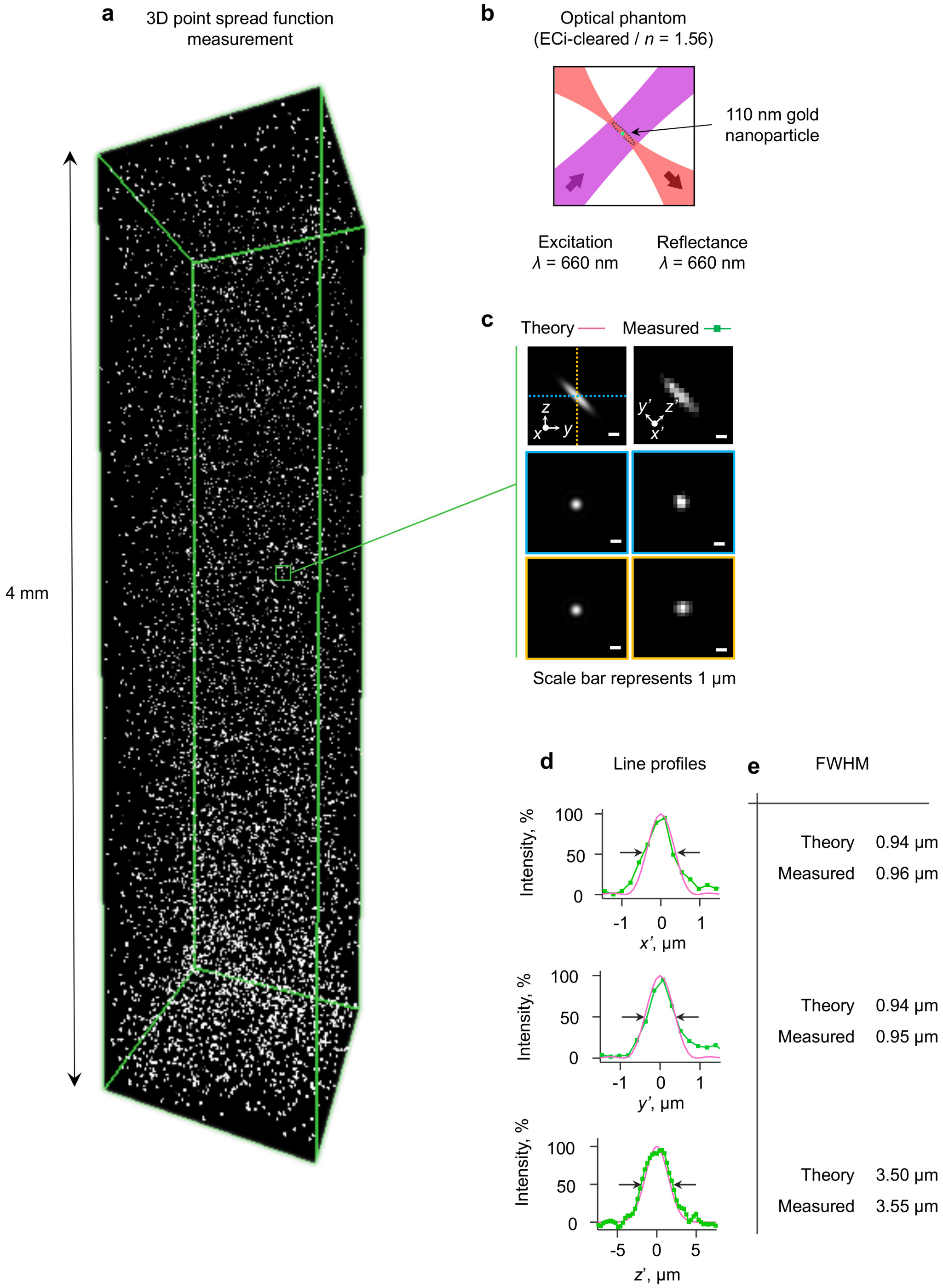


**Supplementary Figure 8. PSF measurement. (a)** 3D rendering of the point spread function measured in an optical phantom to a depth of 4 mm. The optical phantom was comprised with 110 nm gold nanoparticles (1:1000 dilution) in 1% low-melting point agarose. After fabrication, the phantom was dehydrated using ethanol, and cleared using ECi resulting in a refractive index of 1.56. Rather than fluorescence, the reflectance from the gold nanoparticles was measured to assess the point spread function **(c)**. Line profiles and comparisons of the measured point spread function to the theoretical point spread function are shown in **(d)** and **(e)**.

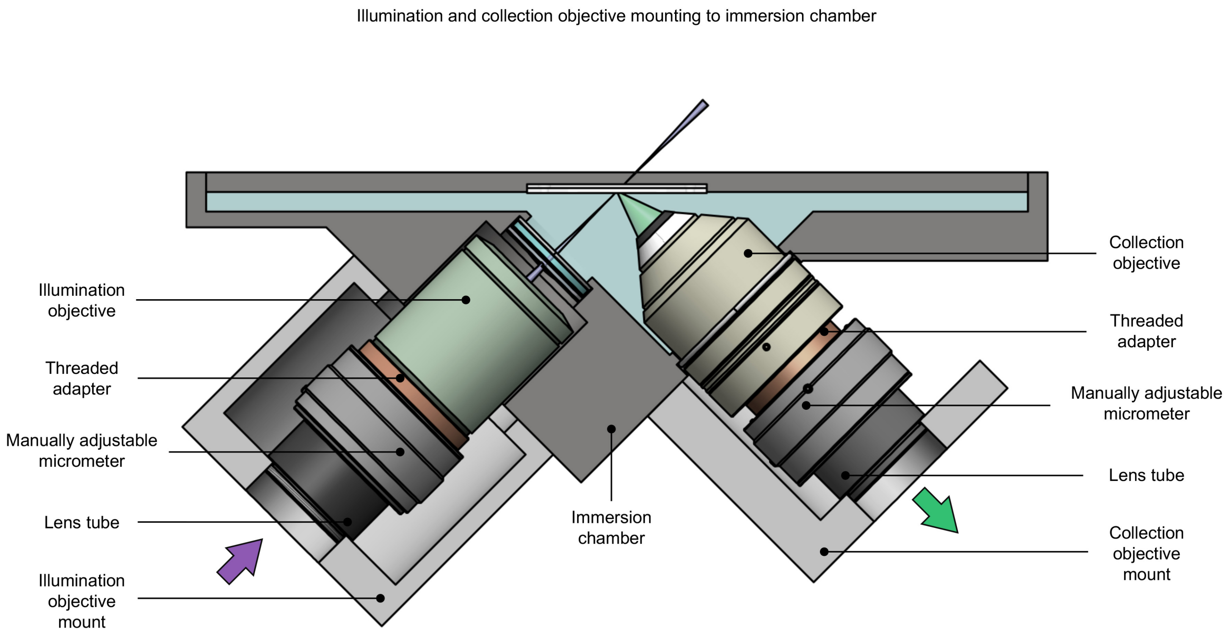


**Supplementary Figure 9. Immersion chamber.** Both the illumination and collection objectives interface with the immersion chamber using custom fabricated objective mounts. The objectives are each attached to the mounts using threaded adapters, manually adjustable micrometers, and lens tubes. The manually adjustable micrometers enable precise adjustment of the objective distances from the beam focus for alignment purposes. The custom components are available as **Supplementary CAD Files**.

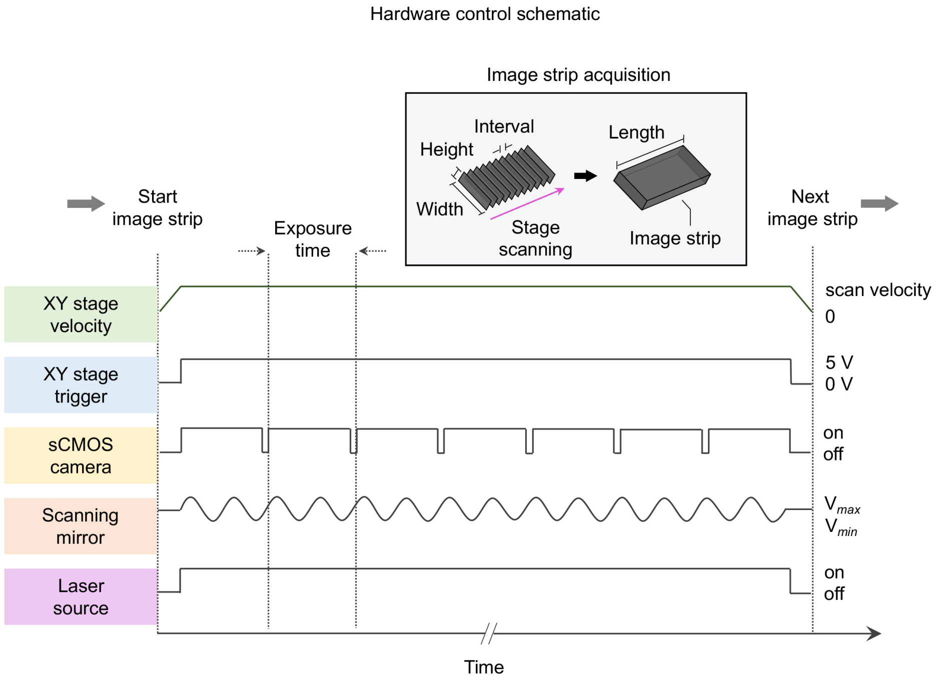


**Supplementary Figure 10. Image strip acquisition.** Image strips are collected with a combination of stage-scanning and lateral/vertical tiling using a motorized XY stage and Z actuators. The stage-scanning firmware is used to send a TTL trigger signal from the XY stage to the sCMOS camera for reproducible start positioning (<1 µm) of each image strip. The spatial interval between successive frames is set to ~0.32 µm, which given the 800 Hz camera framerate, corresponds to a constant stage velocity of ~0.25 mm/sec. The scanning mirror and laser source are activated at the beginning of an image strip, and deactivated at the end of an image strip.

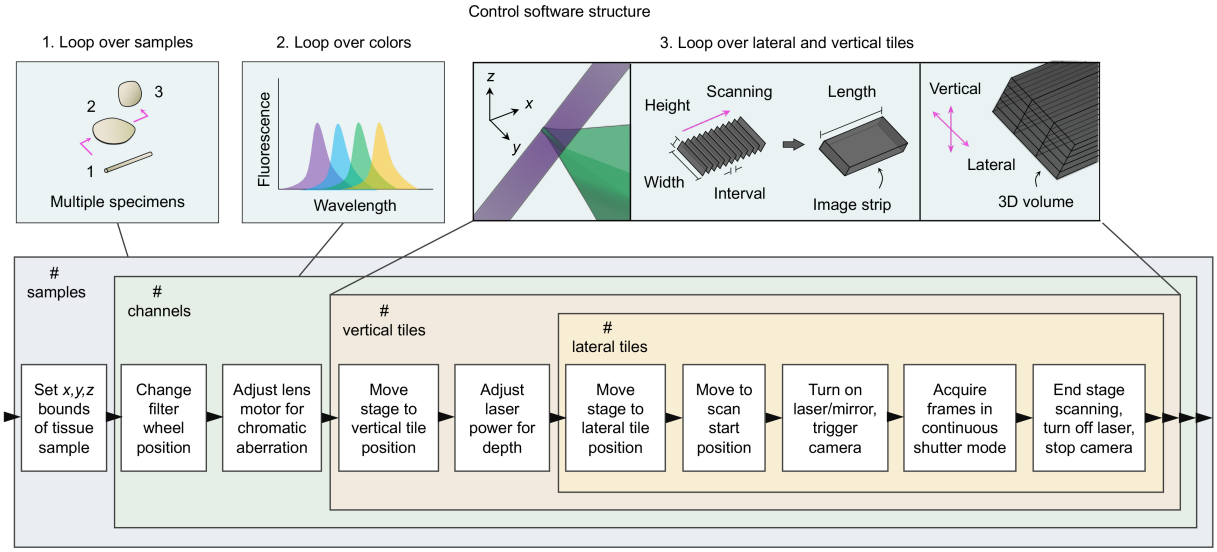


**Supplementary Figure 11. Data acquisition scheme.** Imaging data is collected by using a series of nested loops. The outermost loop scans over the # of samples, with user specification of the *x, y, z* bounds of each tissue sample. The second loop collects the number of user-defined color channels for each sample. Finally, the innermost loops iterate over the vertical and lateral tiles necessary to cover the entire sample.

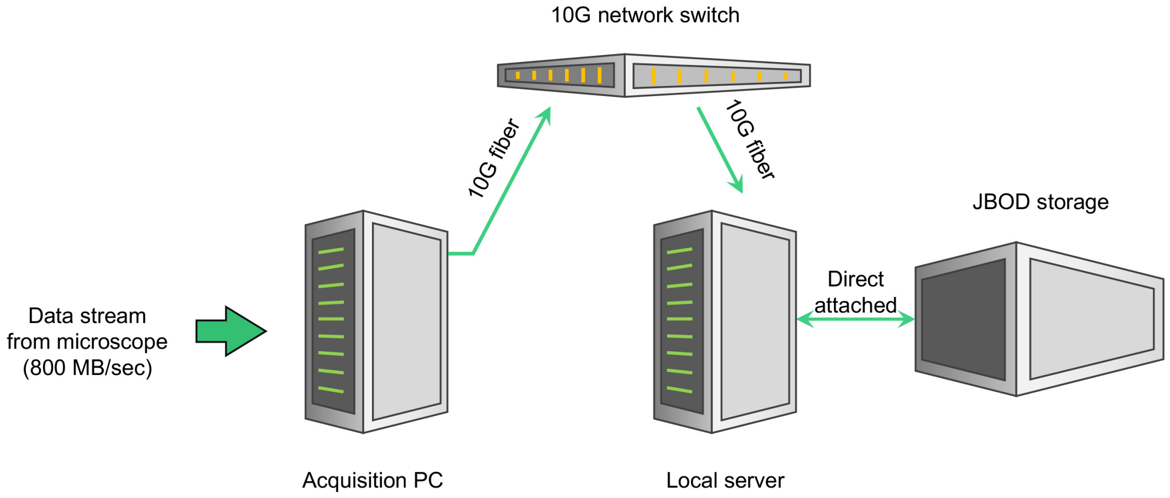


**Supplementary Figure 12. Computing hardware.** In contrast to conventional microscopes, which may acquire data at 1 MB/sec onto a single HDD, with processing by a CPU, and data transfer/storage using 1 Gbit networking to an external drive, light-sheet microscopes acquire data at up to 1 GB/sec. This requires specialized hardware, including a RAID array of SSDs, processing with a GPU, and 10 Gbit transfer to network storage. In our OTLS system, data is streamed to a low specification acquisition PC through a 10Gbit network to a high specification local server with direct-attached JBOD storage. Complete specifications of the system are provided in **Supplementary Table 2**.

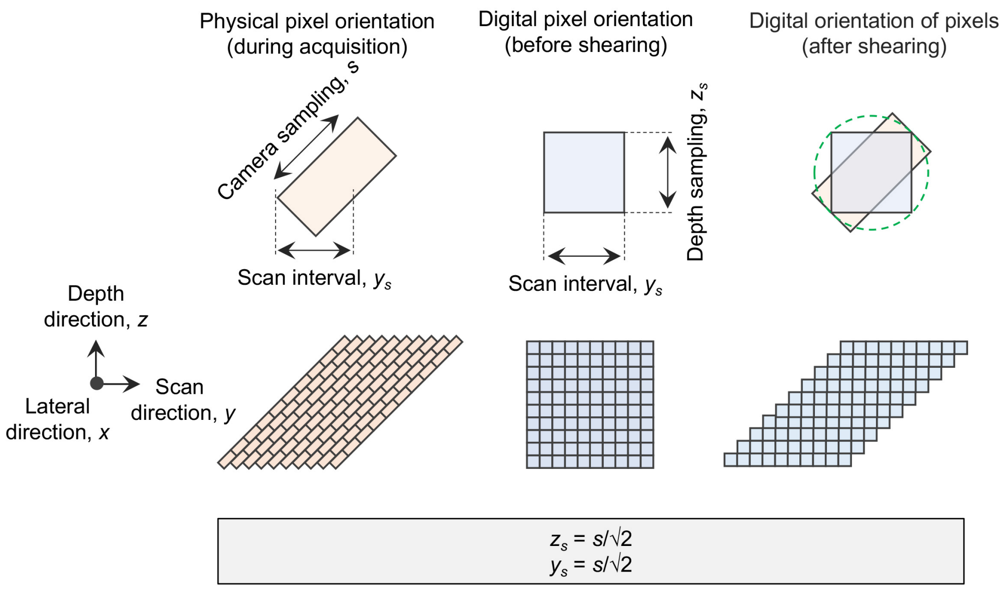


**Supplementary Figure 13. Scan interval between successive frames.** Raw image frames are collected at an oblique 45 deg. angle within samples. These images are initially oriented at 0 deg. in a three-dimensional cube of imaging data. To restore the physical orientation of the image planes, the data must be sheared at the 45 deg. angle. Rather than using an affine transformation calculation, we employ a strategy which shits each row of pixels by an integer pixel offset. For this operation to be valid, the scan interval between successive frames, $y_s$, must be equal to the in-plane camera sampling, *s*, divided by √2**.**

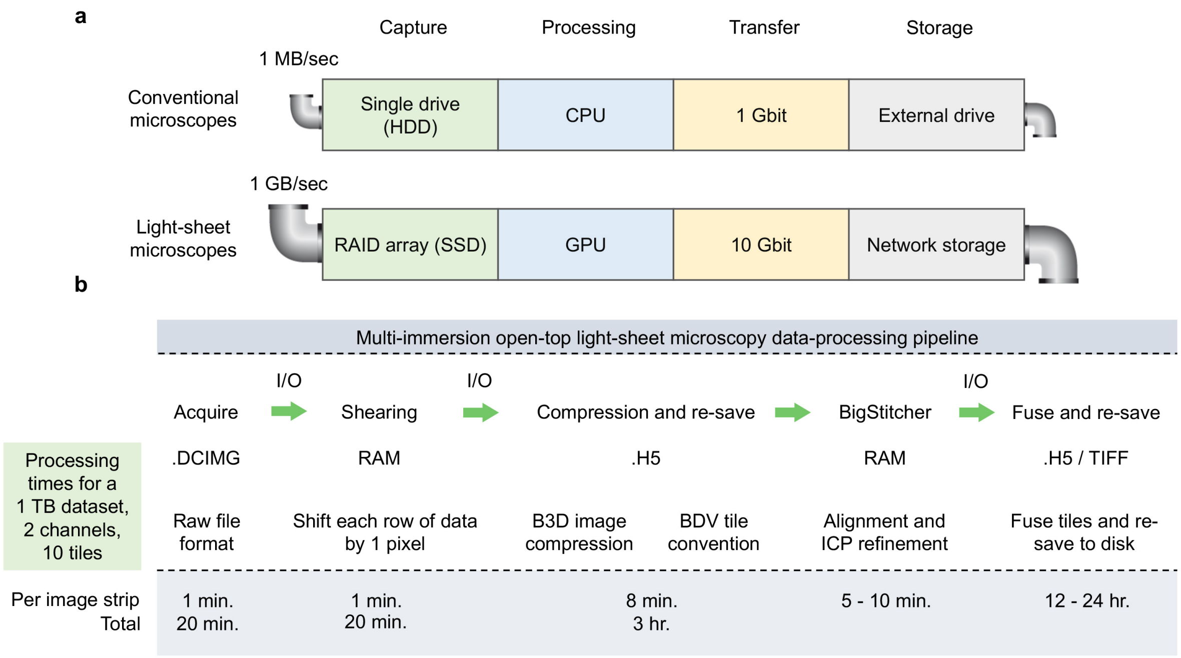


**Supplementary Figure 14. Data processing scheme. (a)** In contrast to conventional microscopes, which may acquire data at 1 MB/sec onto a single HDD, with processing by a CPU, and data transfer/storage using 1 Gbit networking to an external drive, light-sheet microscopes acquire data at up to 1 GB/sec. This requires specialized hardqare, including a RAID array of SSDs, processing with a GPU, and 10 Gbit transfer to network storage. (b) For the multi-immersion OTLS system, each image stripe is stored in a single DCIMG file. These DCIMG files are read into RAM by a DLL compiled using the Hamamatsu DCIMG software development kit (SDK) and first sheared at 45 deg. The data is then written from RAM to disk using the Hierarchical Data Format (HDF5) with the metadata and XML file structured for subsequent analysis using BigStitcher [2]. A custom HDF5 compression filter (B3D) is used with default parameters to provide ~10× compression which is within the noise-limit of the sCMOS camera [3]. This pre-processing routine is applied to all DCIMG files, ultimately resulting in a single HDF5/XML file for BigStitcher. The alignment of all image strips is performed in BigStitcher, and finally fused to disk in either TIFF or HDF5 file formats. The resulting TIFF and HDF5 files are then visualized using open-source and commercial packages, including ImageJ, BigDataViewer, Aivia (DRVision), and Imaris (Bitplane) [4, 5]. Representative processing times for a 1 TB dataset are shown in **(b)**. The processing routines are available as **Supplementary Code Files**. Graphic adapted from [45]

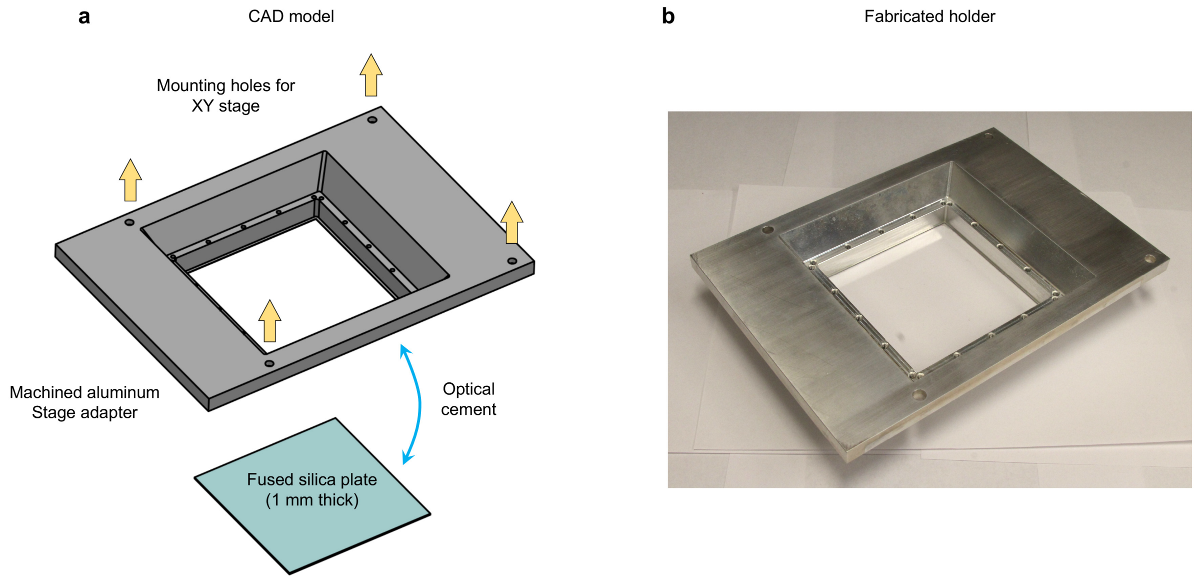


**Supplementary Figure 15. TDE-cleared specimen holder. (a)** CAD rendering of the specimen holder for TDE-clearing. A 10×10 cm by 1 mm thick fused silica plate is optically cemented into a custom machined aluminum adapter which can be mounted to the XY stage. An image of the fabricated holder is shown in **(b).** Files are available as **Supplementary CAD Files**.

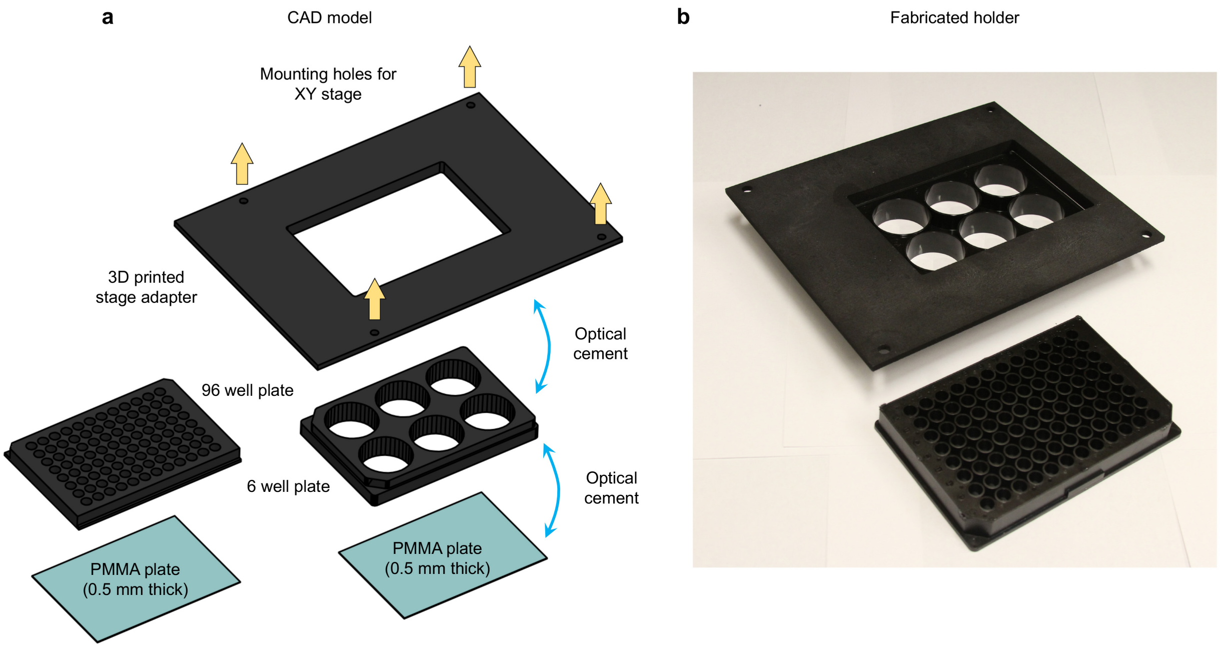


**Supplementary Figure 16. Ce3D-cleared specimen holder. (a)** CAD rendering of the specimen holder for Ce3D-clearing. The bottom surface of plates (with 96 or 6 wells) are retrofitted with a 0.5 mm thick PMMA plate using optical cement. The plates are then cemented to a 3D printed plastic adapter with holes for mounting to the XY stage. An image of the fabricated holder is shown in **(b)**. Files are available as **Supplementary CAD Files**.

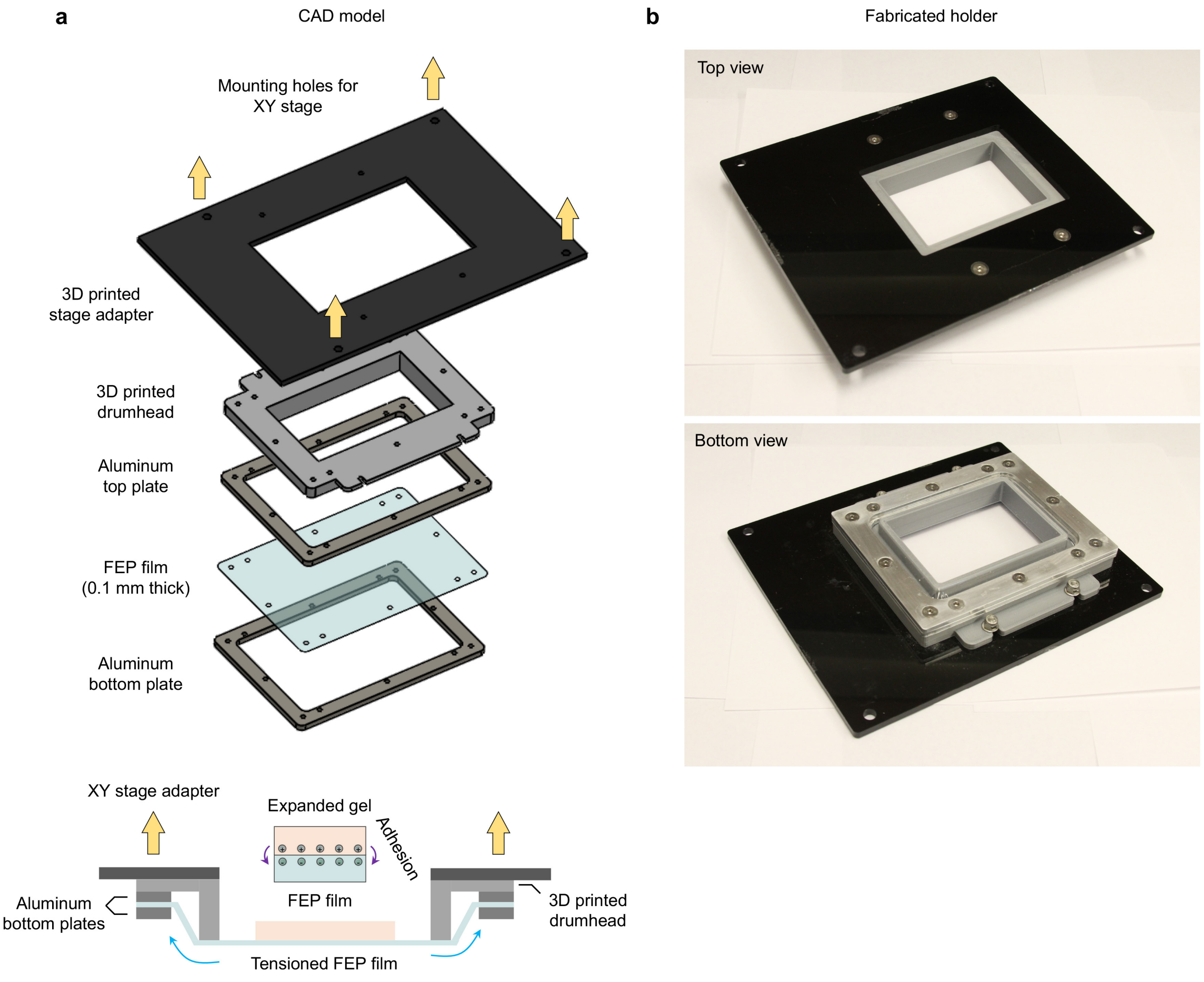


**Supplementary Figure 17. ExM specimen holder. (a)** CAD rendering of the specimen holder for ExM-clearing. A 0.1 mm thick FEP film is held between two custom machined aluminum plates. The FEP film is then tightened over a 3D printed "drumhead". The entire drumhead is then attached to a 3D printed mounting plate which attaches to the XY stage. Prior to imaging, the upper surface of the FEP film is treated with poly-lysine to promote adhesion of the expanded gel to the FEP film and prevent movement of the gel during long imaging sessions. Top and bottom images of the fabricated holder is shown in **(b)**. Files are available as **Supplementary CAD Files**.

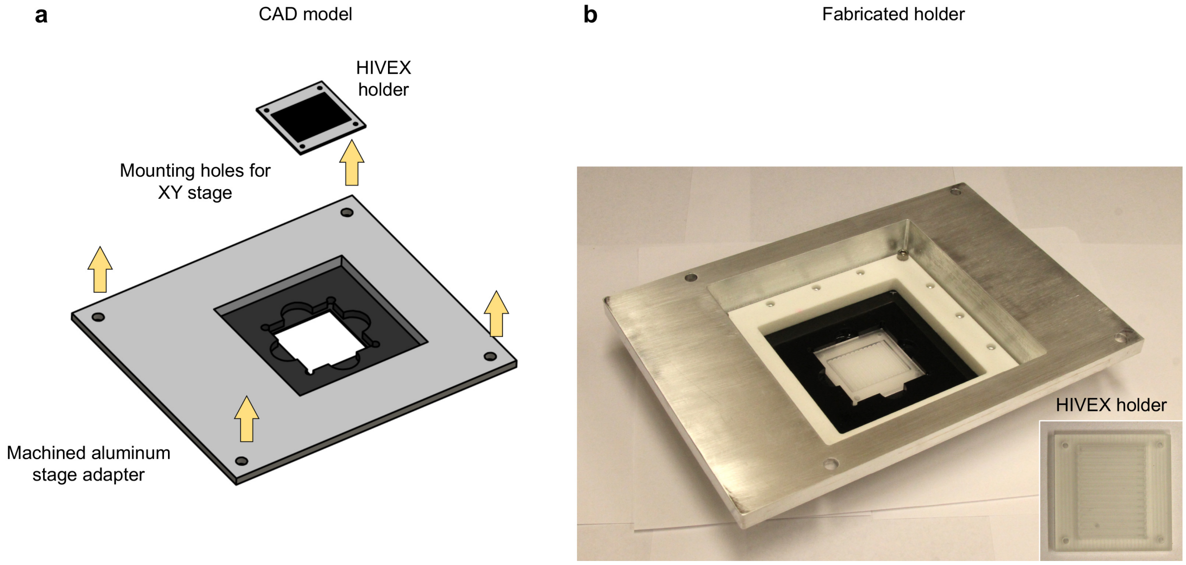


**Supplementary Figure 18. ECi-cleared specimen holder. (a)** CAD rendering of the specimen holder for ECi-clearing. A custom machined HIVEX holder with 13 channels for human biopsies is placed in a custom machined aluminum adapter for mounting to the XY stage. An image of the fabricated holder is shown in **(b)**. Files are available as **Supplementary CAD Files**.

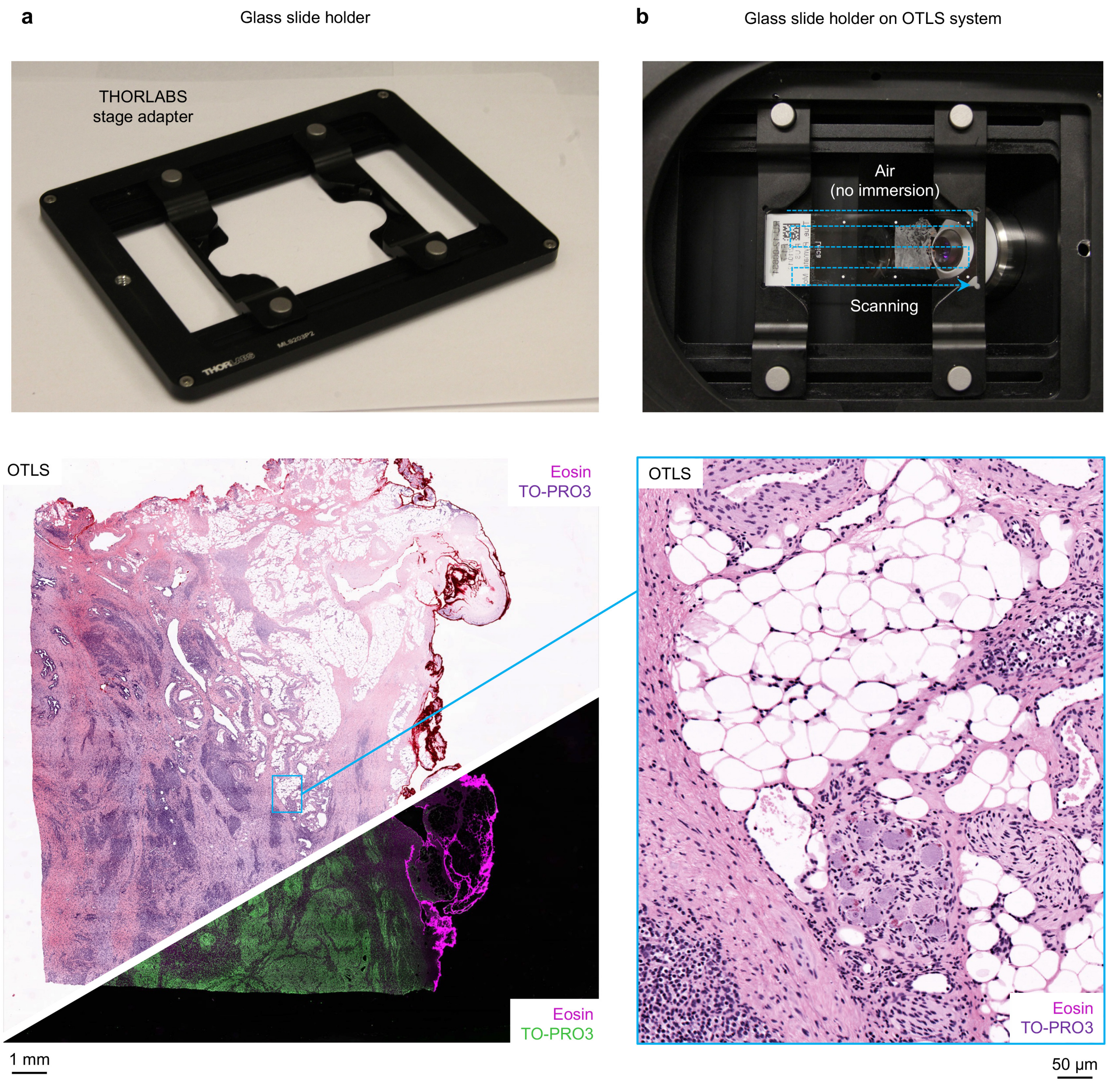


**Supplementary Figure 19. Glass slide holder and whole slide imaging. (a)** THORLABS stage adapter for holding glass slides. **(b)** Representative image of a fluorescently labeled (TO-PRO3 and Eosin) slide-mounted histology section on the OTLS system. Glass slides may be imaged in air (i.e. no immersion) since the tissue sections are thin and optical aberrations only accumulate at larger depths. A whole slide fluorescence image of the slide in **(b)** is shown with pseudo-H&E false coloring (top left) as well as a dark-field false-coloring palette (bottom right) that is more typical for fluorescence microscopy in research settings. A high magnification zoom-in is also shown.

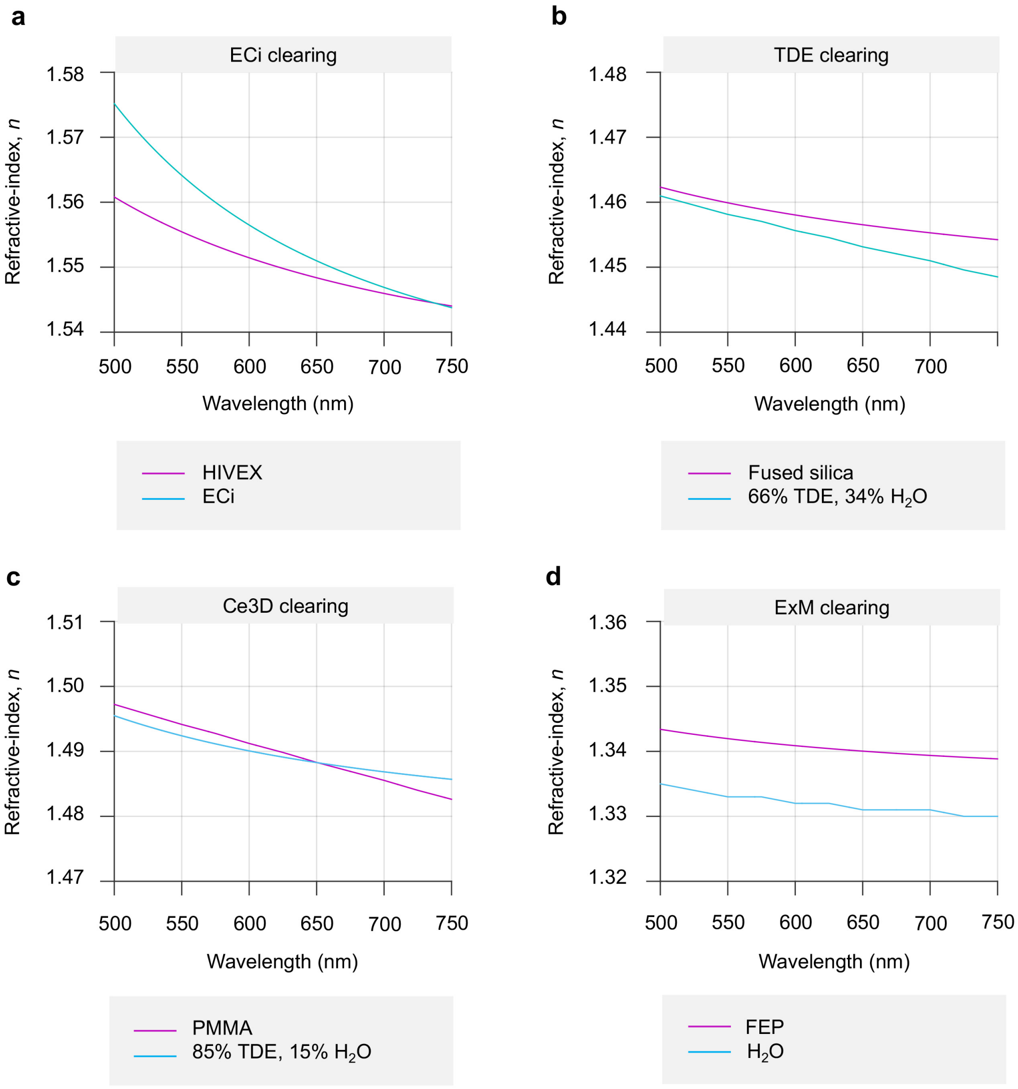


**Supplementary Figure 20. Dispersion curves for specimen holder and clearing reagent combinations. (a-d)** The dispersion curves for ECi clearing (ECi / HIVEX), TDE clearing (66% TDE, 34% $H_2O$ / Fused silica), Ce3D clearing (85% TDE, 15% $H_2O$ / PMMA), and ExM clearing ($H_2O$ / FEP) are shown. Data for materials and reagents were obtained from [38, 46, 47]. Data for the HIVEX (Conant Optical) and FEP (DuPont) materials were obtained from the manufacturers. However, the dispersion of many clearing reagents, particularly multi-compound aqueous-based solutions, are unknown (see **Supplementary Discussion**).

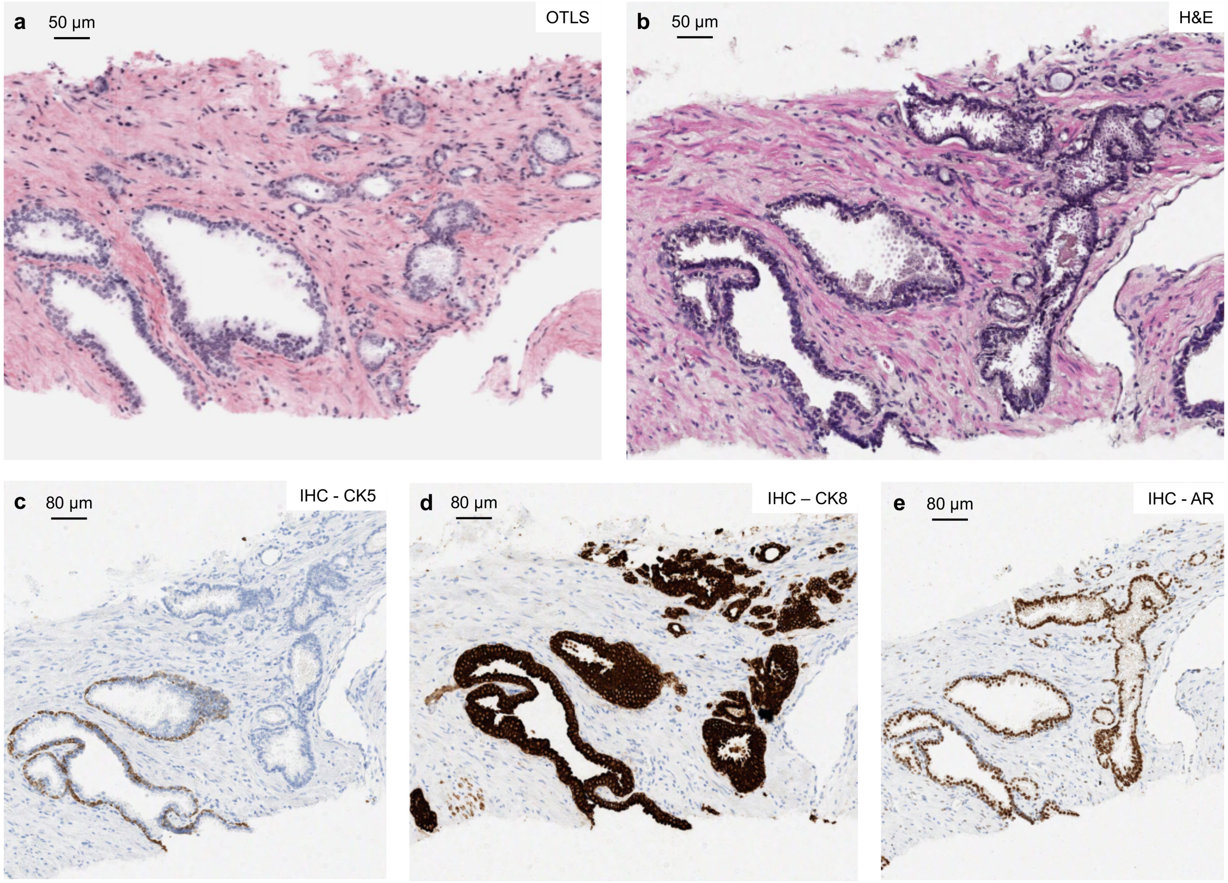


**Supplementary Figure 21. Compatibility of ECi-clearing with downstream histology of human prostate tissue. (a)** 2D OTLS image of an intact human prostate biopsy. **(b-e)** Corresponding H&E and IHC of the same sample and region of interest containing a large vascular channel, benign glands, and well-formed carcinoma glands. Nuclear (AR) and cytoplasmic (CK8, CK5) stains are shown. AR stains the nucleus of all carcinoma and benign glands, CK8 stains the cytoplasm of all carcinoma glands and the luminal epithelium of the benign glands, and CK5 stains the basal cells of the benign glands, but is absent in the carcinoma glands.

**a** Simplified model

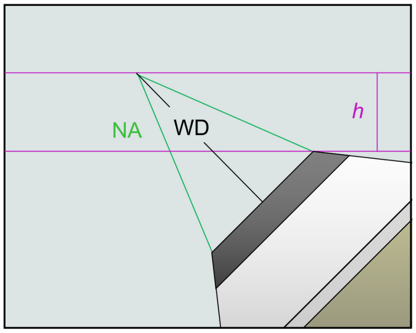


**b** Additional factors considered

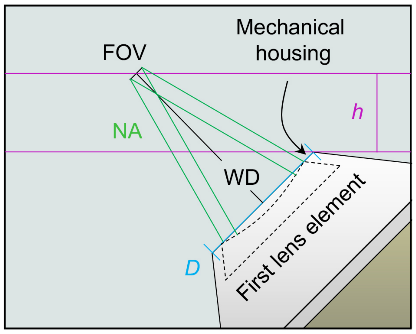


| Variables |
|---|
| *h* – Maximum theoretical imaging depth |
| NA – Numerical aperture |
| D – Mechanical diameter of objective tip |
| WD – Working distance |
| FOV – Field of view |

| Equations |
|---|
| $h$ = WD / √2 x (1– NA) |
| $h_{actual}$ = WD / √2 x (1– sin(tan$^{-1}$(D/(2 xWD)))) |
| BFP diameter = 2 x EFL x NA |

**c** Objectives used in current OTLS system

| Olympus XLFLUOR 340/4X | |
|---|---|
| NA | 0.28 |
| WD | 29.5 mm |
| EFL | 45 mm |
| D | 35 mm |
| FOV | 5.5 mm |
| BFP diameter | 25.2 mm |
| $h$ | ~ 15 mm |
| $h_{actual}$ | ~ 5.7 mm |

| ASI / Special Optics objective | |
|---|---|
| NA | 0.276 |
| WD | 12 mm |
| EFL | 17.4 mm |
| D | 9.8 mm |
| FOV | 1.2 mm |
| BFP diameter | 9.6 mm |
| $h$ | ~ 6.1 mm |
| $h_{actual}$ | ~ 5.1 mm |

**Supplementary Figure 22. Objective design for open-top imaging. (a)** Simplified model depicting an objective oriented at 45 deg. for open-top imaging. The numerical aperture (NA), working distance (WD), and maximum theoretical clearance (*h*), are shown. **(b)** Model with additional factors considered, including the field of view (FOV) and mechanical housing of the objective. The variables and equations for the clearance are shown below. The specifications for both objectives used in the current OTLS system are shown in **(c)**.

### Supplementary Videos

| Name | Thumbnail | Description |
|---|---|---|
| Supplementary Video 1 | 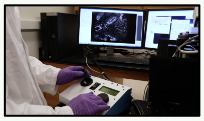 | Video demonstrating the simplicity of mounting samples on the OTLS system, as well as real-time previewing of specimen(s), followed by automated imaging of the specimen(s). |
| Supplementary Video 2 | 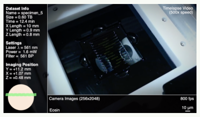 | Time-lapse video demonstrating the operating principles of the OTLS system (image tiling as well as multiple specimens). Shown are 12 ECi-cleared human prostate biopsies, stained with TO-PRO3 (nuclear) and Eosin (cytoplasmic). |
| Supplementary Video 3 | 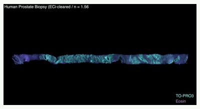 | Surface blend volume rendering of one of the ECi-cleared human prostate biopsies shown in **Fig. 3a**. Color channels are TO-PRO3 (cyan) and Eosin (purple). |
| Supplementary Video 4 | 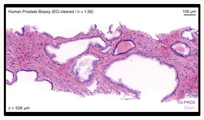 | Z-stack of benign glands in a ECi-cleared human prostate biopsy. The TO-PRO3 and Eosin channels are pseudo-colored to mimic conventional H&E staining. |
| Supplementary Video 5 | 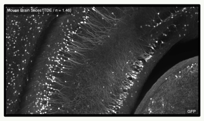 | Maximum intensity volume renderings of 8 TDE-cleared mouse brain slices, imaged successively with the OTLS system. The GFP channel is shown with an inverted grayscale lookup table. |
| Supplementary Video 6 | 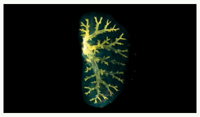 | Maximum intensity volume renderings of the bronchial tree in a Ce3D-cleared mouse lung. Color channels are EpCAM (yellow) and F-actin (blue). |
| Supplementary Video 7 | 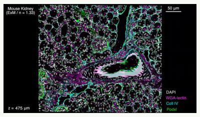 | Z-stack of a small sub-region of an entire ExM-cleared mouse kidney specimen. Color channels are DAPI (white), WGA-lectin (magenta), Coll IV (cyan), and Podxl (green). |
| Supplementary Video 8 | 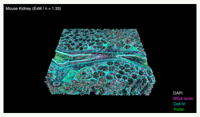 | Surface blend volume rendering of a small sub-region of an entire ExM-cleared mouse kidney specimen |
| Supplementary Video 9 | 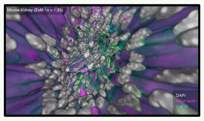 | Fly through of a blood vessel for the expanded mouse kidney shown in Supplementary Videos 8 and 9. |

3D visualizations were generated using Imaris 9.1.2 (Bitplane). Videos were exported from Imaris as raw AVI files with no compression and a resolution of 2560×1440 pixels (2k 16:9). The raw AVIs were then re-saved as AVIs using ImageJ and JPEG compression. The resulting JPEG compressed AVIs were further compressed using Premiere Pro (Adobe) and H.264 compression with a 20 Mbit bitrate. Finally, the supplementary videos were composed using Final Cut Pro X (Apple) and saved as .mp4 files. The timelapse footage in Supplementary Video 1 was captured using a standard webcam (C920, Logitech) controlled using MATLAB R2018b (Mathworks). Supplementary Videos 3 and 4 were false-colored using the Beer-Lambert algorithm [6].

### Supplementary ZEMAX Files

| Name | Description |
| --- | --- |
| **Illumination Optics Files** | |
| ILLUMINATION.ZMX | Parent file containing entire illumination optics model |
| TTN116502-S04_BB.ZMX | TTL-165 Thorlabs tube lens ZEMAX lens file |
| 23522-S03_BB.ZMX | CLS-SL Thorlabs scan lens ZEMAX file |
| **Collection Optics Files** | |
| COLLECTION.ZMX | Parent file containing entire collection optics model |
| TTN116502-S04_BB.ZMX | TTL-165 Thorlabs tube lens ZEMAX lens file |
| 54-10-12@480-910nm reversed_BB.ZMX | Multi-immersion collection objective ZEMAX lens file |

### Supplementary CAD Files

| Name | Description |
| --- | --- |
| **Chamber** | |
| chamber_assembly.SLDASM | Parent file containing entire immersion chamber model |
| 4xobje.SLDASM | Olympus XLFLUOR 340/4X assembly |
| CHAMBER_BLOCK_2_7_2018.SLDPRT | Custom machined immersion chamber block |
| CHAMBER_MOUNT1.SLDPRT | Illumination objective custom chamber mount |
| CHAMBER_MOUNT2.SLDPRT | Collection objective custom chamber mount |
| CLEARING_OBJECTIVE.SLDPRT | Multi-immersion collection objective model |
| LA4725-A.SLDPRT | SIL lens model |
| O_RING.SLDPRT | SIL gasket model |
| O_RING_2.SLDPRT | Collection objective gasket model |
| SM1A24.SLDPRT | Illumination objective threaded adapter |
| SM1A61.SLDPRT | Collection objective threaded adapter |
| SM1L05.SLDPRT | Illumination objective lens tube attachment |
| SM1L05-2.SLDPRT | Collection objective lens tube attachment |
| SM1ZM_1.SLDPRT | Manually adjustable micrometer (part1) |
| SM1ZM_2.SLDPRT | Manually adjustable micrometer (part2) |
| **TDE** | |
| stage_adapter.SLDPRT | Adapter for mounting to XY stage |
| fused_silica_plate.SLDPRT | Fused silica plate |
| **Ce3D** | |
| stage_adapter.SLDPRT | Adapter for mounting to XY stage |
| 6_wellplate.SLDPRT | 6 well plate |
| 96_wellplate.SLDPRT | 96 well plate |
| pmma_plate.SLDPRT | PMMA plate |
| **ExM** | |
| stage_adapter_3.SLDPRT | Adapter for mounting to XY stage |
| plastic_base.SLDPRT | 3D printed drumhead plastic base |
| aluminum_plate.SLDPRT | Aluminum plate (same model for top/bottom plates) |

| | |
|---|---|
| FEP_film.SLDPRT | FEP film |
| **ECi** | |
| stage_adapter.SLDPRT | Adapter for mounting to XY stage |
| hivex_holder.SLDPRT | HIVEX specimen holder for human biopsies |
| **WSI** | |
| MLS203P2.SLDPRT | Thorlabs stage adapter for holding glass slides |

#### Supplementary Code Files

| Name | Description |
|---|---|
| dcimg_hdf5.py | Python file for processing image strips into B3D compressed BDV files.<br><br>Inputs:<br><br>filename: path and name of DCIMG file<br>dir: shearing direction (0 or 1). for this data should be set to 0.<br>idx: index for imaging tile (0 ... # of tiles).<br>binFactor: imaging data binning factor (1, 2, 4, 8). for this data should be set to 1.<br><br>Outputs:<br><br>data.h5: this file will be in the same directory as the input data<br><br>Example usage:<br>>> python dcimg_hdf5.py 000001.dcimg --dir=0 --idx=0 --binFactor=1 |
| bdv_xml.py | Python file for generating the BDV XML file for BigStitcher.<br><br>Inputs:<br>channels: number of color channels<br>tiles: number of imaging tiles in each color channel.<br><br>Outputs:<br>data.xml: this file will be in the same directory as the bdv_xml.py file<br><br>Example usage:<br>>> python bdv_xml.py --channels=1 --tiles=4 |
| DCIMG.dll | DLL file called by dcimg_hdf5.py, compiled using the DCIMG SDK. |

**Supplementary Data** and instructions for installing and testing all of the code files is available at:
https://figshare.com/articles/Supplementary_Data/7685597
