## Supplementary Code Files for "Multi-immersion open-top light-sheet microscope for high-throughput imaging of cleared tissues": bdv_xml.py

```
import argparse
import h5py

if __name__ == '__main__':

	print("Writing BigDataViewer XML file...")

	parser = argparse.ArgumentParser()
	parser.add_argument("--channels", type = int, default = 2, help ="color channels")
	parser.add_argument("--tiles", type = int, default = 10, help ="imaging tiles")
	args = parser.parse_args()

	c = args.channels
	t = args.tiles

	f = open('data.xml', 'w')
	f.write('<?xml version="1.0" encoding="UTF-8"?>\n')
	f.write('<SpimData version="0.2">\n')
	f.write('\t<BasePath type="relative">.</BasePath>\n')
	f.write('\t<SequenceDescription>\n')
	f.write('\t\t<ImageLoader format="bdv.hdf5">\n')
	f.write('\t\t\t<hdf5 type="relative">data.h5</hdf5>\n')
	f.write('\t\t</ImageLoader>\n')
	f.write('\t\t<ViewSetups>\n')

	f2 = h5py.File('data.h5','r')
	dset = f2.get('/t00000/s00/0/cells')
	x = dset.shape[2]
	y = dset.shape[1]
	z = dset.shape[0]

	for i in range (0, c):
		for j in range(0, t):
			f.write('\t\t\t<ViewSetup>\n')
			f.write('\t\t\t\t<id>' + str(t*i+j) + '</id>\n')
			f.write('\t\t\t\t<name>' + str(t*i+j) + '</name>\n')
			f.write('\t\t\t\t<size>' + str(x) + ' ' + str(y) + ' ' + str(z) + '</size>\n')
			f.write('\t\t\t\t<voxelSize>\n')
			f.write('\t\t\t\t\t<unit>um</unit>\n')
			f.write('\t\t\t\t\t<size>0.4455 0.3151 0.3151</size>\n')
			f.write('\t\t\t\t</voxelSize>\n')
			f.write('\t\t\t\t<attributes>\n')
			f.write('\t\t\t\t\t<illumination>0</illumination>\n')
			f.write('\t\t\t\t\t<channel>' + str(i) + '</channel>\n')
			f.write('\t\t\t\t\t<tile>' + str(j) + '</tile>\n')
			f.write('\t\t\t\t\t<angle>0</angle>\n')
			f.write('\t\t\t\t</attributes>\n')
			f.write('\t\t\t</ViewSetup>\n')

	f.write('\t\t\t<Attributes name="illumination">\n')
	f.write('\t\t\t\t<Illumination>\n')
	f.write('\t\t\t\t\t<id>0</id>\n')
	f.write('\t\t\t\t\t<name>0</name>\n')
	f.write('\t\t\t\t</Illumination>\n')
	f.write('\t\t\t</Attributes>\n')
	f.write('\t\t\t<Attributes name="channel">\n')

	for i in range(0, c):
		f.write('\t\t\t\t<Channel>\n')
		f.write('\t\t\t\t\t<id>' + str(i) + '</id>\n')
		f.write('\t\t\t\t\t<name>' + str(i) + '</name>\n')
		f.write('\t\t\t\t</Channel>\n')

	f.write('\t\t\t</Attributes>\n')
	f.write('\t\t\t<Attributes name="tile">\n')

	for i in range(0, t):
		f.write('\t\t\t\t<Tile>\n')
		f.write('\t\t\t\t\t<id>' + str(i) + '</id>\n')
		f.write('\t\t\t\t\t<name>' + str(i) + '</name>\n')
		f.write('\t\t\t\t</Tile>\n')

	f.write('\t\t\t</Attributes>\n')
	f.write('\t\t\t<Attributes name="angle">\n')
	f.write('\t\t\t\t<Illumination>\n')
	f.write('\t\t\t\t\t<id>0</id>\n')
	f.write('\t\t\t\t\t<name>0</name>\n')
	f.write('\t\t\t\t</Illumination>\n')
	f.write('\t\t\t</Attributes>\n')
	f.write('\t\t</ViewSetups>\n')
	f.write('\t\t<Timepoints type="pattern">\n')
	f.write('\t\t\t<integerpattern>0</integerpattern>')
	f.write('\t\t</Timepoints>\n')
	f.write('\t\t<MissingViews />\n')
	f.write('\t</SequenceDescription>\n')

	f.write('\t<ViewRegistrations>\n')
	for i in range(0, c*t):
		f.write('\t\t<ViewRegistration timepoint="0" setup="' + str(i) + '">\n')
		f.write('\t\t\t<ViewTransform type="affine">\n')
		f.write('\t\t\t\t<Name>calibration</Name>\n')
		f.write('\t\t\t\t<affine>1.0 0.0 0.0 0.0 0.0 0.707 0.0 0.0 0.0 0.0 0.707 0.0</affine>\n')
		f.write('\t\t\t</ViewTransform>\n')
		f.write('\t\t</ViewRegistration>')

	f.write('\t</ViewRegistrations>\n')
	f.write('\t<ViewInterestPoints />\n')
	f.write('\t<BoundingBoxes />\n')
	f.write('\t<PointSpreadFunctions />\n')
	f.write('\t<StitchingResults />\n')
	f.write('\t<IntensityAdjustments />\n')
	f.write('</SpimData>')
```
