## Supplementary Code Files for "Multi-immersion open-top light-sheet microscope for high-throughput imaging of cleared tissues": dcimg_hdf5.py

```
import argparse
import numpy as np, ctypes
import os.path
import timeit
import gc
import warnings
import scipy.ndimage
import h5py
from tqdm import tqdm

def shear_images(img_3d, direction, height, nFrame, binFactor):

	print('Shearing images...')

	start = timeit.default_timer()			
	
	for i in tqdm(xrange(height)):
		if direction:
			img_3d[i,i:nFrame,:] = np.clip(img_3d[i,height:img_3d.shape[1]-i,:],100*binFactor*binFactor, 65535) - 100*binFactor*binFactor

		else:
			img_3d[i,height-1-i:nFrame,:] = np.clip(img_3d[i,0:img_3d.shape[1]-height+1+i,:],100*binFactor*binFactor, 65535) - 100*binFactor*binFactor

	if direction:
		img_3d = img_3d[:,0:nFrame-height,:]
	else:
		img_3d = img_3d[:,height:nFrame,:]

	stop = timeit.default_timer()
	print('Shearing time: {0}s'.format(stop-start))

	return img_3d

def save_images(img_3d, idx, binFactor, save_dir):
	
	print('Writing images...')

	start = timeit.default_timer()

	res_list = (1, 2, 4, 8)

	res_np = np.zeros((len(res_list), 3), dtype = 'float64')
	res_np[:,0] = res_list
	res_np[:,1] = res_list
	res_np[:,2] = res_list

	dest = save_dir + '\\data.h5'
	f = h5py.File(dest,'a')
	
	sgroup = f.create_group('/s' + str(idx).zfill(2))
	resolutions = f.require_dataset('/s' + str(idx).zfill(2) + '/resolutions', chunks = (res_np.shape), dtype = 'float64', shape = (res_np.shape), data = res_np)

	subdiv_np = np.zeros((len(res_list), 3), dtype = 'uint32')

	if idx == 0:
		tgroup = f.create_group('/t00000')
	
	f.close()

	for z in range(len(res_list)-1, -1, -1):

		chunkSize1 = 32/binFactor
		chunkSize2 = 256
		chunkSize3 = 256/binFactor
			
		f = h5py.File(dest,'a')
		res = res_list[z]

		subdiv_np[z, 0] = chunkSize1
		subdiv_np[z, 1] = chunkSize2
		subdiv_np[z, 2] = chunkSize3

		resgroup = f.create_group('/t00000/s' + str(idx).zfill(2) + '/' + str(z))

		if z == 0:

			print('Writing block level ' + str(z))

			data = f.require_dataset('/t00000/s' + str(idx).zfill(2) + '/' + str(z) + '/cells', chunks = (chunkSize1, chunkSize2, chunkSize3), dtype = 'int16', shape = img_3d.shape, compression = 32016, compression_opts=(round(2*1000), 1, round(2.1845*1000), 0, round(1.6*1000)))			
			
			chunks = np.arange(0, img_3d.shape[0], chunkSize1, dtype = 'int')
			chunks = np.append(chunks, img_3d.shape[0])

			for i in range(1,len(chunks)):

				print('Writing chunk ' + str(i) + '/' + str(len(chunks)-1))
				N1 = chunks[i-1]
				N2 = chunks[i]
				data[N1:N2] = img_3d[N1:N2]
				f.flush()
				gc.collect()
		else:

			print('Writing block level ' + str(z))

			with warnings.catch_warnings():
				warnings.simplefilter("ignore")
				img_3d_temp = scipy.ndimage.interpolation.zoom(img_3d, float(1.0/res), order = 1, mode = 'nearest')

			data = f.require_dataset('/t00000/s' + str(idx).zfill(2) + '/' + str(z) + '/cells', chunks = (chunkSize1, chunkSize2, chunkSize3), dtype = 'int16', shape = img_3d_temp.shape, compression = 32016, compression_opts=(round(2*1000), 1, round(2.1845*1000), 0, round(1.6*1000)))			
			
			chunks = np.arange(0, img_3d_temp.shape[0], chunkSize1, dtype = 'int')
			chunks = np.append(chunks, img_3d_temp.shape[0])

			for i in range(1,len(chunks)):

				print('Writing chunk ' + str(i) + '/' + str(len(chunks)-1))
				N1 = chunks[i-1]
				N2 = chunks[i]
				data[N1:N2] = img_3d_temp[N1:N2]
				f.flush()
				gc.collect()
	
	f = h5py.File(dest,'a')
	subdivisions = f.require_dataset('/s' + str(idx).zfill(2) + '/subdivisions', chunks = (res_np.shape), dtype = 'uint32', shape = (subdiv_np.shape), data = subdiv_np)
	f.close()

	stop = timeit.default_timer()
	print('Writing time: {0}s'.format(stop-start))

if __name__ == '__main__':

	print("Pre-processing code")

	global_start = timeit.default_timer()

	tqdm.monitor_interval = 0

	parser = argparse.ArgumentParser()
	parser.add_argument("filename", help='file name of .dcimg')
	parser.add_argument("--dir", choices=range(2), type = int, default = 0, help="direction")
	parser.add_argument("--idx", type = int, default = 0, help ="channel/tile number")
	parser.add_argument("--binFactor", type = int, default = 1, help ="camera binning")
	args = parser.parse_args()

	filename = args.filename
	direction = args.dir
	idx = args.idx
	binFactor = args.binFactor

	# paths
	cwd = os.getcwd()
	file_path = os.path.abspath(filename)
	lib_path='.\\DCIMG.dll'
	dir_name = os.path.dirname(filename)
	slash=dir_name.find('\\')
	dir_name=dir_name[slash:]
	save_dir = os.path.abspath(dir_name)

	# check the nesassary files
	if os.path.exists(file_path):
		print('File found.')
	else:
		print('File not found.')
		sys.exit()
		
	if os.path.exists(lib_path):
		print('Dynamic-link library found.')
	else:
		print('Dynamic-link library not found.')
		sys.exit()
	# load the DLL to read the .DCIMG file
	libc = ctypes.cdll.LoadLibrary(lib_path)
	if libc.OpenDcimg(file_path):
		print('Dynamic-link library loaded.')
	else:
		print('Dynamic-link library not loaded.')
		sys.exit()

	# get the sessioin number of the dcimg file
	nTotalSession = libc.GetTotalSessionCount()
	session_index = 0

	nFrame = ctypes.c_int()
	width = ctypes.c_int()
	height = ctypes.c_int()
	rowbyte = ctypes.c_int()

	start = timeit.default_timer()

	if not libc.GetImageInformation(session_index,ctypes.byref(nFrame),
							 ctypes.byref(width),ctypes.byref(height),ctypes.byref(rowbyte)):
		print('Failed to get information about the image.')
		sys.exit()
	else:
		nFrame, width, height, rowbyte = nFrame.value, width.value, height.value,rowbyte.value
		# raw is the raw data of the image
		raw_type=(ctypes.c_ubyte*(rowbyte* height))
		raw = raw_type()
		
		print('Session #{0}: {1} frames.'.format(session_index, nFrame))
		print('Width x height: {0} x {1}.'.format(width, height))

		#read images
		print('Reading images...')
		img_3d = np.zeros((nFrame, height, width), dtype = 'uint16')
		for frame_index in tqdm(xrange(nFrame)): 
			libc.AccessRawFrame(frame_index, session_index,ctypes.byref(raw))
			img_3d[frame_index] = np.frombuffer(np.ctypeslib.as_array(raw), dtype = '<u2').reshape(height,width)

		stop = timeit.default_timer()
		print('Reading time: {0}s'.format(stop-start))

		img_3d = img_3d.transpose((1,0,2))

		img_3d = shear_images(img_3d, direction, height, nFrame, binFactor)
			
		img_3d = img_3d.transpose((1,0,2))
		img_3d = np.flip(img_3d,1)
		img_3d = img_3d.transpose((1,0,2))

		save_images(img_3d, idx, binFactor, save_dir)

	global_stop = timeit.default_timer()
	print('Total time: {0}s'.format(global_stop-global_start))

	libc.CloseDcimg()
```
